# Endogenously expressed P2X5 includes a proportion of isoforms or transcript variants sensitive to extracellular ATP

**DOI:** 10.1101/2025.01.10.632374

**Authors:** Xiao-Na Yang, Dong-Ping Wang, Chen-Hui Shen, Xi-Chen, Hui Yang, Xing Zhou, Chang-Run Guo, Yun Tian, Michael X. Zhu, Ye Yu

**Author notes:** equal contributions to this work (Xiao-Na Yang, Dong-Ping Wang, Chen-Hui Shen, Xi-Chen). Correspondence and requests for materials should be addressed to Y.Y.

## Abstract

The P2X receptor family comprises ATP-gated non-selective cation channels central to physiological processes across the nervous, immune, cardiovascular, respiratory, and reproductive systems. While P2X1, P2X2, P2X3, P2X4, and P2X7 are well-established as ATP-sensitive, P2X5 isoforms and transcript variants (TVs) have traditionally been considered ATP-insensitive, limiting their functional exploration. This study identifies previously overlooked ATP-sensitive P2X5 isoforms across diverse species. Through gene profiling and next-generation sequencing (NGS), we found that the zfP2X5^TV2^ isoform dominates ATP-sensitive forms in zebrafish. In mice, ATP-sensitive mP2X5^G317^ (mP2X5.1) comprises ∼90% of transcripts, while in rats, exon 3-containing rP2X5^F191^ accounts for over 70%. In human cell lines, ATP-sensitive isoforms retaining exons 3, 7, and 10 represent ∼15-30% of P2X5 transcripts. RNA-seq from human tissues confirms frequent retention of exons 3 and 7 and polymorphic exon 10 expression. ATP-sensitive P2X5 variants were also identified in chickens, bullfrogs, dogs, cows, and naked mole-rats. These findings challenge the prevailing assumption of ATP-insensitivity, highlighting the need to reassess P2X5’s roles in physiological and pathological contexts.

## Introduction

The P2X receptor family encompasses ATP-gated, non-selective cation channels that mediate NaL, KL, and Ca²L permeability (Roberts, Vial et al., 2006, Schmid & Evans, 2019), with certain subtypes also demonstrating ClL conductance (Bo, Jiang et al., 2003, Browne, Cao et al., 2011, Schiller, Jacobson et al., 2022). To date, seven subtypes (P2X1-7) have been cloned, all of which assemble into functional trimers that may be either homomeric or heteromeric (Kopp, Krautloher et al., 2019, Nicke, 2008, Saul, Hausmann et al., 2013). All subtypes, except P2X6, can form functional homomeric trimers (Schmid & Evans, 2019), while all, excluding the controversial P2X7 subtype (Kopp et al., 2019, Nicke, 2008), are also capable of forming heteromeric complexes (Saul et al., 2013). Key subtypes, including P2X1, P2X2, P2X3, P2X4, and P2X7, are central to sensory functions such as pain perception, taste, hypoxia detection, and bladder distension. They also mediate neural transmission in the central and peripheral nervous systems, immune and inflammatory responses, cardiovascular regulation, and renal filtration (Coddou, Yan et al., 2011b, Illes, Muller et al., 2021, King, 2023). P2X5, first cloned from celiac ganglia, exhibits expression in regions like the trigeminal mesencephalic nucleus, heart, and spinal cord (Schmid & Evans, 2019). Further studies demonstrated its broader distribution in the immune system, heart, and skeletal muscle, linking it to processes such as inflammatory bone loss, immune regulation, cancer metastasis, cell proliferation and differentiation, and polycystic kidney disease (Coddou et al., 2011b, Illes et al., 2021, King, 2023, Schmid & Evans, 2019). Despite its diverse expression and potential functional significance, P2X5 has remained underexplored. This is likely due to early findings suggesting that homomeric P2X5 trimers have minimal or no ATP sensitivity (Coddou, Stojilkovic et al., 2011a, King, 2023, Schmid & Evans, 2019). Additionally, P2X5 has been proposed to function primarily in heteromeric complexes with P2X1, forming P2X1/5 assemblies (Coddou et al., 2011b, King, 2023, Lalo, Pankratov et al., 2008, Schmid & Evans, 2019). However, this model remains contested, necessitating further detailed investigation (King, 2023).

When first cloned, the human, mouse, and rat P2X5 receptors (hP2X5, mP2X5, and rP2X5) displayed minimal ATP-induced currents (Collo, North et al., 1996, Cox, Barmina et al., 2001, Le, Paquet et al., 1997). For mP2X5 and rP2X5, these weak responses were linked to subtle amino acid substitutions despite the presence of complete coding sequences (King, 2023, Sun, Liu et al., 2019). In humans, most hP2X5 variants lack exon 10, the critical region encoding the second transmembrane domain (TM2), thereby rendering the receptor non-functional (Kotnis, Bingham et al., 2010). Smita et al. uncovered that reinstating exon 10 fully restores TM2 and rescues hP2X5 functionality (Kotnis et al., 2010). However, such functional variants are exceedingly rare, having been identified in only one African American individual among a cohort of 46 samples representing diverse ethnic groups (Kotnis et al., 2010). Notably, functional ATP-sensitive P2X5 isoforms are more frequently observed in non-mammalian species (Jensik, Holbird et al., 2001, Low, Kuwada et al., 2008, Meyer, Groschel-Stewart et al., 1999, Ruppelt, Ma et al., 2001), as summarized in Table EV1. In zebrafish, two splice variants, TV1 and TV2, exhibit opposite ATP sensitivities: TV1, found in somite tissue, is non-functional, whereas TV2, detected in muscle tissue, is highly ATP-sensitive (Kucenas, Li et al., 2003, Low et al., 2008). Thus, while P2X5 homomeric trimers are generally considered non-functional, particularly in common mammalian, they have neither been disappeared during evolution nor rendered pseudogenes like certain ion channels such as TRPC2 (King, 2023). The remarkable diversity of P2X5 transcripts and their ATP sensitivities underscores a complex regulatory system yet to be fully understood (King, 2023). With advances in molecular and sequencing technologies, deeper insights into P2X5 functions in various tissues, developmental stages, and physiological or pathological contexts are anticipated, necessitating a reexamination of its functional roles (Hattori & Gouaux, 2012, King, 2023, Schmid & Evans, 2019, Stark, Grzelak et al., 2019).

To unravel the mechanisms underlying ATP responsiveness in P2X5 receptor splice variants and isoforms, we conducted a comprehensive analysis across multiple species. Using the zebrafish P2X5 (zfP2X5) variants TV1 and TV2 as models, we identified key regions that contribute to the functional differences between these variants through mutation screening, electrophysiological assays, and molecular dynamics (MD) simulations. While these regions display partial amino acid conservation, they are crucial for both P2X gating and ATP recognition, suggesting that mutations or single nucleotide polymorphisms (SNPs) in these areas could influence ATP responsiveness in a tissue- and development-specific manner. Through T-A cloning and next generation sequencing (NGS) of zebrafish *P2RX5* clones, we found that the functional zfP2X5^TV2^ variant is consistently expressed across all developmental stages, maintaining significant ATP responsiveness despite the presence of high-frequency mutations. In mammalian models, our mRNA analysis identified a strong correlation between rat mP2X5^G317^ and mouse rP2X5^F191^ with strong ATP responses. Similarly, T-A cloning and NGS from human airway epithelial cells (16HBE) and gastric epithelial cells (GES-1) revealed that exons 3, 7, and 10 are retained in about 15-30% of transcripts, a crucial factor for sustaining strong ATP responsiveness in hP2X5. RNA-seq analysis from the NCBI human database revealed that exons 3 and 7 are expressed at high levels, whereas exon 10 shows lower expression in certain organs, suggesting polymorphic variations in the *P2RX5* gene. Furthermore, plasmid-based ATP-induced current assessments confirmed that full-length P2X5 receptors from other species, such as chickens, cows, dog, and naked mole rats, exhibit strong ATP responses, further supporting the involvement of P2X5 in diverse physiological and pathological processes. These results challenge the assumption that P2X5 trimers are weak ATP responders, revealing that isoforms with stronger responsiveness may be more prevalent across tissues and developmental stages than previously thought. This underscores the need for further study into their physiological and pathological significance.

## Results

### Divergent ATP responsiveness of zebrafish P2X5 splice variants arises from non-conserved amino acids involved in channel gating, ATP recognition or surface expressions

Functional investigations of the P2X5 receptor have uncovered a dichotomy in ATP responsiveness between mammalian and non-mammalian species. In mammals, including rats, mice, and humans, P2X5 typically exists in a weak or non-functional form (Collo et al., 1996, Cox et al., 2001, Le et al., 1997), whereas in non-mammalian species such as zebrafish and frogs, functional phenotypes characterized by robust ATP responsiveness (Jensik et al., 2001, Low et al., 2008, Meyer et al., 1999) (Table EV1). Interestingly, the zebrafish *P2RX5* gene encodes two splice variants, zfP2X5^TV^and zfP2X5^TV2^, which exhibit distinctly opposite ATP responsiveness, with zfP2X5^TV1^ being non-functional and zfP2X5^TV2^ exhibiting functional characteristics (Kucenas et al., 2003, Low et al., 2008). The functional divergence between these two splice variants offers valuable insights into the potential mechanisms underlying the transition between different genetic forms of P2X5. The sequence alignment of the zfP2X5^TV1^ and zfP2X5^TV2^ variants revealed nine amino acid differences, located at residues 39, 116, 144, and within the 283-288 loop region (Fig 1A, and ref. (Low et al., 2008)). Mutations corresponding to the non-functional zfP2X5^TV1^ variant were introduced into the functional zfP2X5^TV2^, specificallyzfP2X5^TV2-V39G^, zfP2X5^TV2-Y116H^, zfP2X5^TV2-H144L^, and zfP2X5^TV2-283-288(SQHSVT-YTALGP)^, leading to a complete loss of ATP responsiveness at all these sites (Fig. 1B). Notably, zfP2X5^TV2-V39G^ and zfP2X5^TV2-Y116H^ mutants exhibited a complete loss of membrane expression (Fig. 1C), suggesting that these mutations interfere with protein folding or prevent proper trafficking to the membrane. In contrast, the zfP2X5^TV2-H144L^ and zfP2X5^TV2-283-288(SQHSVT-YTALGP)^ mutants retained normal membrane expression (Fig 1C). Both mutations lie close to the ATP-binding pocket, with residues 283-288 located in the left flipper domain (Figs. 1D and EV1A) and H144 located in the head domain (Fig. EV3A, B). These regions play a crucial role in ATP binding-induced conformational changes and may also be involved in ATP recognition (Coddou et al., 2011a, Wang, Sun et al., 2017, Zhao, Wang et al., 2014), exhibiting considerable amino acid divergence across the P2X1-7 subtypes and between different species of the same subtype (Kawate, Michel et al., 2009) (Fig. EV1A and EV3A). Additionally, to pinpoint the key amino acids responsible for the functional loss inzfP2X5^TV2-283-288(SQHSVT-YTALGP)^, each of the six amino acids in this region was substituted with the corresponding amino acids from zfP2X5^TV1^. The resulting mutants, zfP2X5^TV2-S283Y^ and zfP2X5^TV2-Q284T^ exhibited significantly reduced ATP responsiveness, while the double mutant, zfP2X5^TV2-S283Y/Q284T^ almost entirely abolished ATP-induced currents (current densities = 11.48 ± 1.5, 4.57 ± 3.7, and 1.24 ± 0.5 pA/pF, respectively; P < 0.05 vs. 134.39 ± 30.4 pA/pF in zfP2X5^TV2-WT^, n = 3-8, one-way ANOVA followed by Dunnett’s multiple comparison test, Fig. EV1B,C). All mutants exhibited normal membrane expression (Fig. EV1D).

**Fig. 1.**
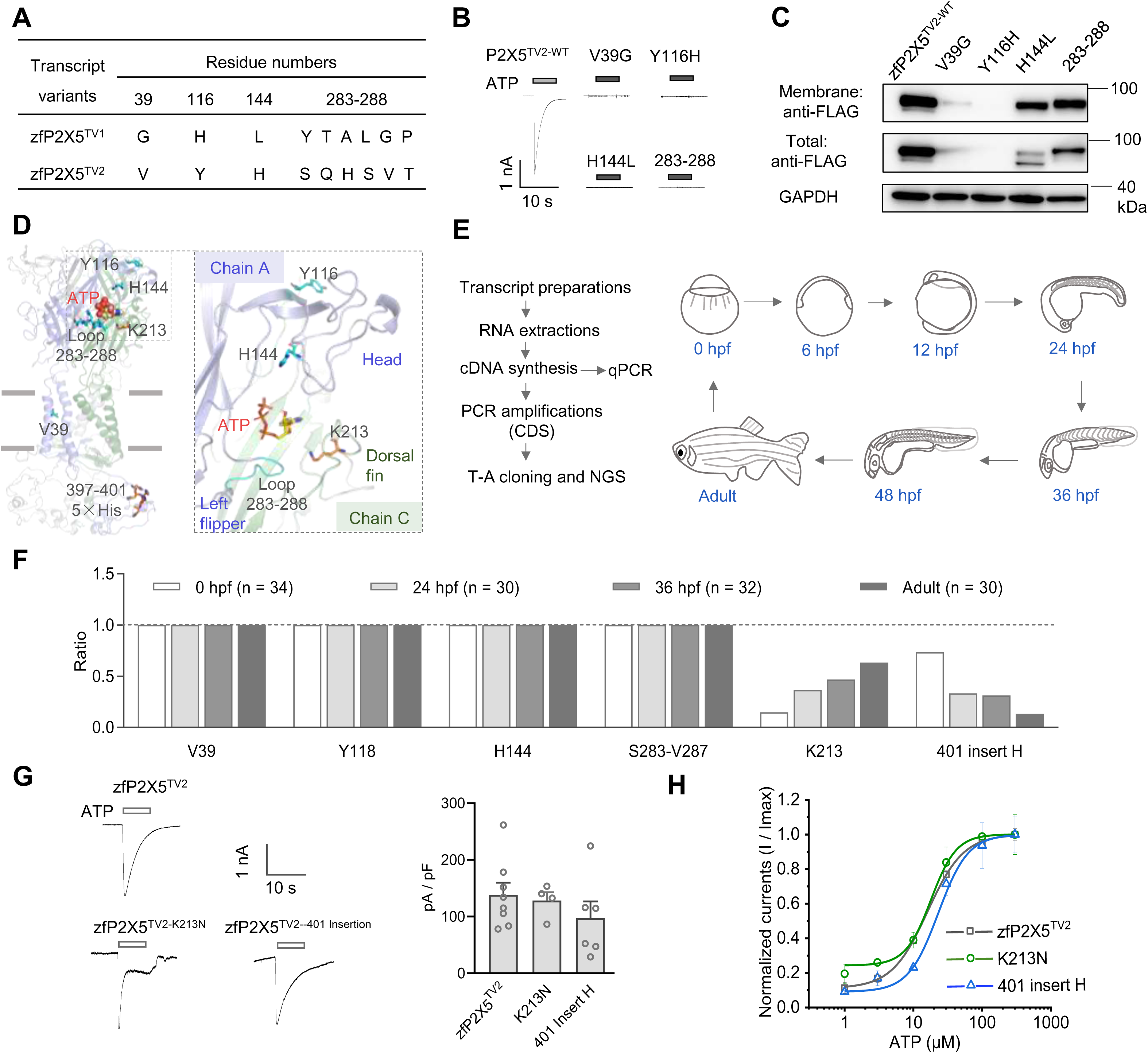
ATP responsiveness of zebrafish P2X5 splice variants diverges due to non-conserved amino acids in gating, ATP recognition, or surface expression. **(A)** Amino acid differences between two zebrafish P2X5 splice variants. **(B-C)** Representative current traces in response to ATP (B) and Western blot analysis of the total and surface protein expressions (C) for zfP2X5 WT and its mutants. **(D)** Homology model of zfP2X5^-TV2^ in the open state, highlighting amino acid positions that differ from those in zfP2X5^-TV1^. **(E)** Schematic representation of zebrafish sampling at various developmental stages. **(F)** Statistical analysis of P2X5 cloning results across different developmental stages of zebrafish. **(G)** Representative current traces (left panel) and pooled data (right panel) for zfP2X5 and its mutants (ATP: 100 μM). Data are expressed as mean ± SEM (n = 4-8). P > 0.05 versus WT, one-way ANOVA followed by Dunnett’s multiple comparisons test, F(2, 15) = 0.84. **(H)** Concentration-response curves for ATP in zfP2X5 WT and its mutants. Data are expressed as mean ± SEM (n = 3).

The homology models of zfP2X5^TV2^, built on the *apo* and open crystal structures of zfP2X4 (PDB IDs: 4DW0, and 4DW1), indicates that the residues S283 and Q284 are located within the left flipper domain. This placement suggests that they may facilitate relative movement between the left flipper and dorsal fin, an event analogous to the gating mechanism observed in zfP2X4 (Wang et al., 2017, Zhao et al., 2014). Such movement is hypothesized to be critical for channel activation (Fig. EV1E, and ref. (Sun et al., 2019). To explore this hypothesis further, we performed conventional molecular dynamics (CMD) simulations using the zfP2X5^TV2-S283Y^ mutant. The substitution of Ser283 with Tyr results in the repositioning of the phenolic hydroxyl group of Tyr to a position closer to the low body domain (Fig. EV2A). This structural alteration leads to a significant shift in the fluctuation range of the adenine nitrogen bond angle of ATP in the CMD simulations, as compared to the zfP2X5^TV2-WT^ (Fig. EV2E). Moreover, this mutation significantly altered the interactions between ATP and key residues, including K68, K70, Y186, and K188 (Fig. EV2B, F). Specifically, the interaction with Y186 was enhanced via water bridging, whereas the hydrophobic interactions were reduced (Fig. EV2F). These changes triggered spatial rearrangements in the positions of residues Y283, Y186, F199, and F190. Notably, the aromatic rings of Y283 and F199 formed π-π interactions, and the aromatic rings of Y186 and F190 created a domino-like effect, collectively influencing ATP binding to the receptor (Fig. EV2C-D).

In response to the steric hindrance introduced by Tyr at position 283, we investigated the possibility of restoring the conformational integrity and ATP responsiveness of this region by reducing the steric bulk of neighboring residues. Remarkably, the mutants zfP2X5^TV2-S283Y/Y186F^, zfP2X5^TV2-S283Y/F190A^ and zfP2X5^TV2-S283Y/F199A^ led to significant recovery of ATP responsiveness, which had been compromised in the S283Y mutant (current densities = 258 ± 23, 128 ± 4, and 148 ± 9 pA/pF, respectively, compared to 24.4 ± 3.3 pA/pF for S283Y; P < 0.05, n = 3-8, one-way ANOVA followed by Dunnett’s multiple comparison test, Fig. EV2G). Additionally, macro-path analysis revealed that 10 μM ATP was sufficient to activate multiple channels in zfP2X5^TV2^, whereas the zfP2X5^TV2-S283Y^ mutant required a higher concentration of ATP to activate a marginally greater number of channels (Fig. EV2H). Both mutants displayed stable single-channel conductance in the 8-10 pS, suggesting that the S283Y substitution modifies ATP interactions with the orthosteric binding pocket (Fig. EV2A, E).

Residue H144, located within the non-conserved region of the head domain (Fig. EV3A), undergoes a pronounced conformational shift from the apo to the open state (Fig. EV3B). The zfP2X5^TV2-H144L^ mutant, which corresponds to the residue found in the TV1 variant, shows no response to ATP, whereas the naturally occurring histidine in the TV2 variant elicits a significant ATP-induced current (16.1 ± 7.8 vs. 171.9 ± 15.9 pA/pF, P < 0.01, n = 5-24, one-way ANOVA followed by Dunnett’s multiple comparison test, Fig. EV3B, C). While differences in glycosylation between the head domain and neighboring glycosylation sites were noted (Hattori & Gouaux, 2012), no significant variation in membrane expression levels was observed among the mutants (P > 0.05, one-way ANOVA followed by Dunnett’s multiple comparison test, n = 3-26, Fig. EV3D, E). These observations suggest that the H144L mutation does not substantially affect protein folding or trafficking. Saturation mutagenesis at this position identified a range of mutations that resulted in diminished or abolished ATP responsiveness (P < 0.05 vs. zfP2X5^TV2^, one-way ANOVA followed by Dunnett’s multiple comparison test, Fig. EV3C). Further membrane expression analysis of several of these mutants, including H144E, H144D, and H144I, demonstrated that the observed functional impairments, including reduced ATP responsiveness, were independent of membrane expression levels or glycosylation status (Fig. EV3D, E). These findings indicate that the gating defect in the zfP2X5^TV2-H144L^ mutant is likely the primary determinant of its loss of function.

These data suggest that the variation in ATP responsiveness between the zebrafish P2X5 transcript variants, zfP2X5^TV1^ and zfP2X5^TV2^, is predominantly driven by mutations in non-conserved residues within the gating and ATP recognition regions of the channel. Moreover, this finding implies that the *zfP2RX5* gene may exhibit tissue-, developmental stage-, or pathology-dependent regulation of ATP responsiveness through specific mutations in hypervariable regions, thereby contributing to differential physiological or pathological functions.

### In the AB strain of zebrafish, P2X5 exhibits strong ATP responsiveness at various developmental stages, with zfP2X5^TV2^ emerging as the predominant variant

Despite we provide evidence delineating the strong/weak ATP response profiles of zfP2X5^TV1^ and zfP2X5^TV2^, prior sequencing studies (Appelbaum, Skariah et al., 2007, Kucenas et al., 2003, Low et al., 2008) failed to provide a comprehensive understanding of the tissue-wide distribution of these variants. Furthermore, it remains unclear whether functional and non-functional variants coexist, or whether their relative expression fluctuates at different stages of development. To address these gaps, we performed an extensive analysis of the *zfP2RX5* gene sequence in the AB-line zebrafish and evaluated the functional properties of the corresponding isoforms or TVs. We first extracted mRNA from whole embryos and adult zebrafish at various developmental stages (0-, 6-, 12-, 24-, 36-, and 48-hours post-fertilization (hpf)) (Fig.1E). Quantitative PCR analysis revealed that P2X5 expression was low at early developmental stages (0, 6, and 12 hpf), but increased significantly at 24, 36, and 48 hpf, before declining again in adult fish (513.7 ± 33.8, 512.4 ± 55.9 and 152.1 ± 7.9, for 24 hpf, 36 hpf and adult, respectively, n = 4; Fig. EV4A).

Sequencing of positive clones obtained via T-A cloning demonstrated that zfP2X5^TV2^ was the sole transcript expressed at all stages, with no detectable presence of zfP2X5^TV1^ (Figs. 1F and EV4B). Interestingly, clones from all stages exhibited mutations at two key positions: the K213 residue in the dorsal fin domain (Fig. 1D) was mutated to asparagine (K213N) with a mutation frequency of 92%, and a histidine insertion was observed at position 401 in the intracellular domain (Fig. 1D), with a mutation probability of 96% (Fig. 1F). To eliminate the possibility of missing low-expression variants due to limited cloning efficiency, we selected three representative developmental stages (0 hpf, 24 hpf, and adult) that displayed a “low-high-low” expression models of P2X5 for high-throughput next generation sequencing (NGS). The sequencing results corroborated our cloning findings, revealing that zfP2X5^TV2^ was the predominant variant at all stages, with two consistent mutations: K213N and a histidine insertion at position 401. Other mutations, each with a frequency of less than 3%, were cataloged (Table EV2).

To assess the functional consequences of the K213N mutation and histidine insertion at position 401, we generated corresponding mutant constructs and evaluated their ATP-induced currents. The results demonstrated that neither mutation impaired ATP responsiveness: the K213N mutation caused a slight reduction in plateau current compared to zfP2X5^TV2^, but the peak current densities were not significantly different between the mutants (128.2 ± 14.6 and 97.1 ± 29.5 pA/pF vs. 138.5 ± 21.2 pA/pF, respectively, P > 0.05, n = 4-8, one-way ANOVA followed by Dunnett’s multiple comparisons; Fig. 1G). Moreover, the apparent ATP affinities (EC_50_, the concentration of ATP yielding half maximum currents) for zfP2X5^TV2-K213N^, zfP2X5^TV2-401-His^ insertion, and zfP2X^5WT^ were similar (16.1 ± 5.4, 18.0 ± 2.3, and 23.0 ± 10.3 μM, respectively, Fig. 1G, H). These findings indicate that, across developmental stages, P2X5 in zebrafish predominantly functions as a highly ATP-responsive receptor.

### In C57BL/6 mice, P2X5 exhibits strong ATP responsiveness, with mP2X5.1^G317^ being the predominant variant

While the initially cloned zfP2X5^TV1^ was characterized by a lack of ATP response (Kucenas et al., 2003), our analysis of zebrafish AB strain tissues at various developmental stages revealed that all P2X5 variants, particularly the zfP2X5^TV2^, are ATP-responsive (Fig.1). This prompted us to explore whether other mammalian P2X5 isoforms, previously thought to be non-functional, might also demonstrate functionality. To investigate this, we examined P2X5 mRNA expression in both murine and human tissues/cells and assessed the functional properties of the corresponding receptors. Previous studies have revealed that mP2X5 exists as three distinct 3’-transcript variants (Cox et al., 2001): mP2X5.1, mP2X5.2, and mP2X5.3 (denoted as m5.1, etc., Fig. 2A, B). The earliest cloned wild-type P2X5 (mP2X5-WT) differs from m5.1 at a single nucleotide in position 317 within the β14-fold region close to the ATP-binding pocket, with mP2X5-WT containing arginine (Arg) and m5.1 containing glycine (Gly) at this position (Fig. 2B,C). Interestingly, the three isoforms listed in the NCBI and Uniport databases also feature Gly at this critical position. Electrophysiological analysis confirmed that the P2X5 variant with Arg at position 317 (mP2X5-WT^R317^) displayed weak ATP responsiveness, while the m5.1, m5.2, and m5.3 variants, all containing Gly at this position, exhibited strong ATP responses, with no significant differences in current densities (41.2 ± 5.3, 45.7 ± 7.1, and 39.2 ± 3.2 pA/pF, respectively, P > 0.05, n = 5-8, one-way ANOVA followed by Dunnett’s multiple comparisons, Fig. 2D,E). These results indicate that 3’-alternative splicing does not impair P2X5 functionality, but the specific residue at position 317, whether Arg or Gly, plays a decisive role in determining ATP responsiveness.

**Fig. 2.**
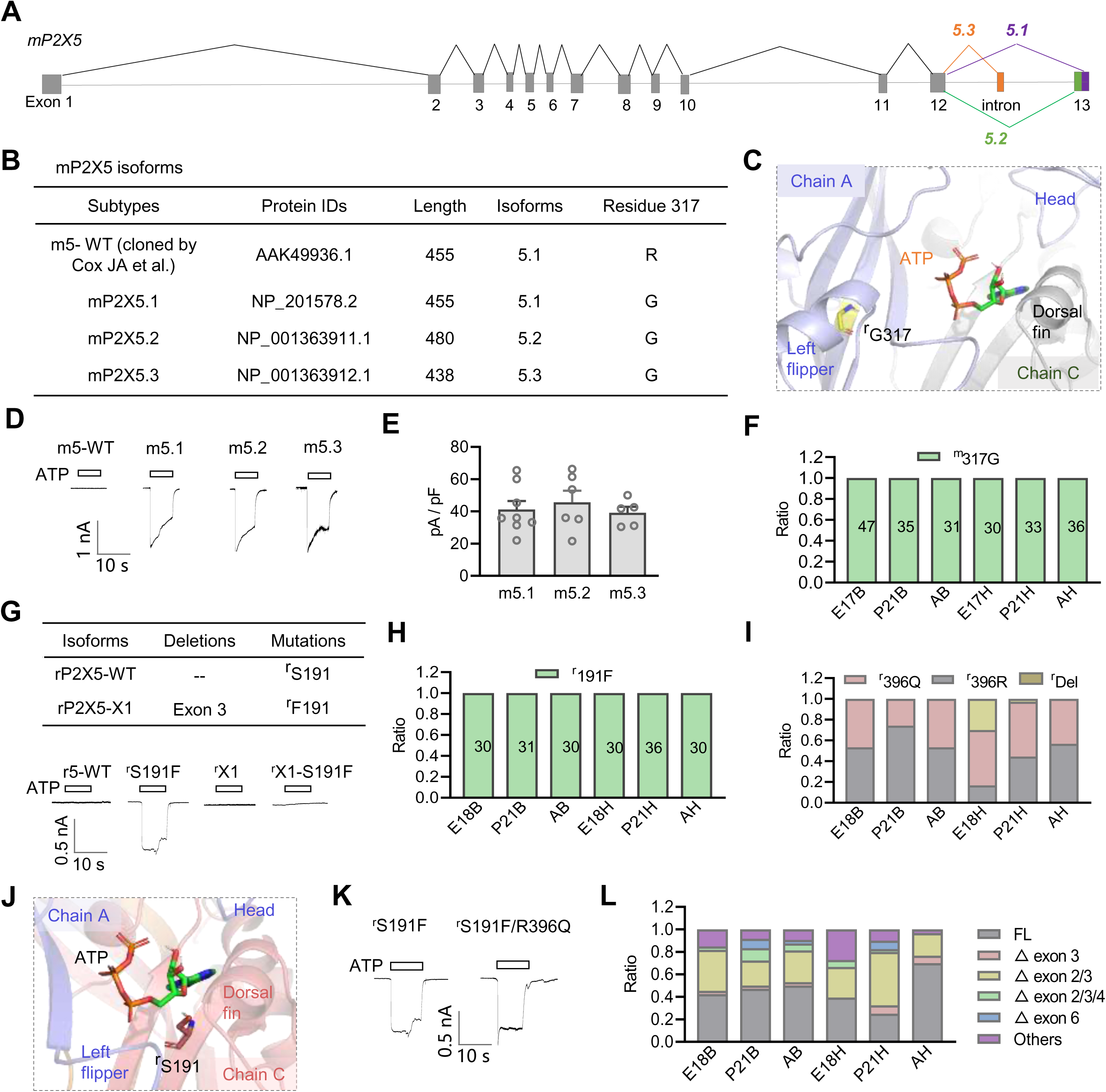
Functional and genetic expressions of P2X5 receptors in mouse and rat. **(A)** Exon distribution patterns of different transcripts of m*P2X5*. (**B**) Sequence characteristics of the mP2X5 isoforms. (**C**) Homology model of mP2X5-ATP complex (using 4DW1 as the template) showing the position of the G317. (**D-E**) Representative current traces (D) and pooled data (E) for mP2X5 and its mutants (ATP: 300 μM, n = 5-8). Data are expressed as mean ± SEM One-way ANOVA with Tukey’s post hoc test, F(2, 16) = 0.31, P = 0.30, P > 0.05. (**F)** Statistical analysis of the ratio of Gly317 in P2X5 receptors across various developmental stages in mouse tissues. E17B, P21B, AB, E17H, P21H, and AH represent the brain and heart from 17-day-old embryonic mice, 21-day-old postnatal mice, and adult mice, respectively. The numbers in the bars represent T-A clone quantities (n ≥ 30). (**G**) Sequence characteristics of rP2X5 isoforms (upper panel) and representative current traces (ATP: 300 μM, lower panel). (**H-I)** Statistical analysis of the ratio of Phe191 (H) and Glu396/Arg396 (I) in P2X5 receptors across developmental stages in rat tissues, determined by cloning of the P2RX5 gene. E18B, P21B, AB, E18H, P21H, and AH represent the brain and heart from 18-day-old embryonic rats, 21-day-old postnatal rats, and adult rats, respectively. (**J)** Homology model of mP2X5-ATP complex showing the position of the S191 residue. (**K)** Representative current traces for rP2X5 S191F and S191F/R396Q mutants. (**L)** Statistical analysis of exon deletion ratios in P2RX5 gene expression in rat tissues at different developmental stages.

To further investigate P2X5 expression patterns in C57BL/6 mice, we cloned and analyzed mP2X5 from heart and brain tissues at three stages: embryonic day 17 (E17), postnatal day 21 (P21), and adulthood (Fig. EV5A). All clones at position 317 contained Gly (Fig. 2F). Notably, the N-terminal sequences detected in these clones revealed the presence of mP2X5.2, with expression levels highest at E17B (19.1%) and lowest at P21B (0%) (Fig. EV5B and Table EV3). No mP2X5.3 sequence was detected (Fig. EV5B and Table EV3). Additionally, we identified novel isoforms, including insertions in intron 2 or intron 4, exon 2-3 deletions, and partial deletions in exon 7 (Table EV4).

To ensure comprehensive data analysis, particularly at position 317, we pooled the mP2X5 clones and performed high-throughput NGS (Fig. EV5C). As expected, sequencing revealed that over 99% of the mP2X5 receptors expressed Gly at position 317, with brain and heart tissues showing 99.7% Gly at this position (Fig. EV5D), and no high-frequency nonsynonymous mutations were detected at other positions. Furthermore, the N-terminal spliced form 5.1 accounted for more than 90% of the total sequence (92.4% in brain and 91.7% in heart, Fig. EV5D). These results suggest that murine mP2X5 is a highly functional ATP receptor, contrasting with previous studies (Collo et al., 1996, Cox et al., 2001, King, 2023).

### In Sprague-Dawley (SD) rats, the rP2X5 receptor is predominantly ATP-responsive

Initially cloned by Collo G and colleagues (Collo et al., 1996), the rP2X5 receptor (rP2X5^WT^) contains a serine residue at position 191, which results in an ATP non-responsive phenotype (Fig. 2G, ref.(Collo et al., 1996)). In addition to this canonical sequence, the most recent full-length rP2X5 sequence, denoted rP2X5^S191F^ (Collo et al., 1996), and an isoform lacking exon 3 (P2X5^X1^) (Garcia-Guzman, Soto et al., 1996), both exhibit phenylalanine at position 191. To further investigate the functional characteristics of these variants, we constructed plasmids encoding the rP2X5^S191F^ and P2X5^X1^ isoforms and expressed them in HEK-293 cells. Electrophysiological analyses revealed that rP2X5^S191F^ exhibited a strong ATP response, whereas both the exon 3-deleted P2X5^X1^ and the P2X5X1^S191F^ variants failed to respond to ATP (Fig. 2G). The residue at position 191 is situated within the ATP-binding pocket, a critical site for receptor function (Fig. 2J). The enhanced ATP responsiveness of rP2X5^S191F^ may be attributed to the augmentation of synergistic interactions between the left flipper and dorsal fin domains, a mechanism previously detailed in our earlier studies (Burnstock & Verkhratsky, 2010).

To ascertain whether rP2X5 expressed *in vivo* is ATP-responsive, we performed gene expression analyses and cloning of rP2X5 from heart and brain tissues at three developmental stages: embryonic day 18 (E18), postnatal day 21 (P21), and adulthood. Higher levels of receptor expression were detected in the adult brain and heart tissues (n = 4, Fig. EV5F), with all clones derived from the three time points containing phenylalanine at position 191 (n = 30–36, Fig. 2H). Notably, a high-frequency mutation at position 396 (R396Q) was identified (Fig. 2I), although it did not impact the functional response of rP2X5 (59.9 ± 16.3 and 87.4 ± 15.7 pA/pF for S191F vs. S191F/R396Q, respectively, P > 0.05, n = 4, Student’s t-test, Figs. 2K and EV5G).

The full-length rP2X5 sequence was consistently expressed in the brain at over 40% across all stages (46.7%, 54.8%, and 53.3% for E18, P21, and adult, respectively, Fig. 2L and Table EV5), with similar phenomena observed in the heart, where the full-length form was more prominently expressed, especially in adulthood (70.0% for adults, Fig. 2L). At earlier stages, such as P21H, the expression was lower, ranging from 20 to 30% (Fig. 2L). The exon 3-deleted P2X5^X1^ isoform was detected, alongside novel isoforms, including those with exon 6 deletions (Fig. 2L). Notably, deletions in exons 2/3 or 2/3/4 led to frameshift mutations (Table EV6).

High-throughput NGS of pooled clones from the three time points revealed that position 191 was consistently occupied by phenylalanine (99.6% in both brain and heart, Fig. EV5E), with a substantial proportion of transcripts containing exon 3 (78.7% and 76.1% in brain and heart, respectively; Fig. EV5E). These data suggest that, at least in SD rats, the full-length P2X5 receptor is highly functional and is expressed at significantly higher levels than other isoforms.

### Normal human cell lines express a subset of hP2X5 receptors capable of exhibiting a robust response to ATP

We further explored the existence of ATP-responsive isoforms of the human P2X5 receptor. So far, only one tissue sample from an African American individual has been found to harbor the full-length exon 10 sequence of the hP2X5 receptor (hP2X5-FL, Fig. 3A), which demonstrates a strong ATP response (Kotnis et al., 2010). The NCBI database catalogues five distinct isoforms of the hP2X5 receptor, differentiated by the presence or absence of exons 3 and 10, as well as specific splice variations within exon 7, notably a modification at the K205-S206 site (Fig. 3B). Isoform A, often referred to as hP2X5-WT, lacks exon 10 within the transmembrane (TM) region, resulting in a loss of ATP responsiveness. Other isoforms exhibit variations such as exon 3 deletions and alterations in exon 7 splice sites, which affect the receptor’s ATP-binding pocket in the dorsal fin region (Fig.3D). Additionally, a selective splicing event at the 3’ end of exon 3 (A3SS, Alternative 3’ Splice Site) leads to a substitution from KS to N (Fig. 3A-D). To investigate the functional consequences of these isoforms, we generated constructs for each isoform and expressed them in HEK-293 cells. Electrophysiological measurements revealed that exon skipping events in exons 3 and 10, as well as A3SS at exon 7, led to a complete loss of ATP responsiveness in the hP2X5-FL receptor (Fig. 3C). These findings underscore the critical importance of the proper expression of exons 3, 7, and 10 for the functional ATP responsiveness of hP2X5 receptors.

**Fig. 3.**
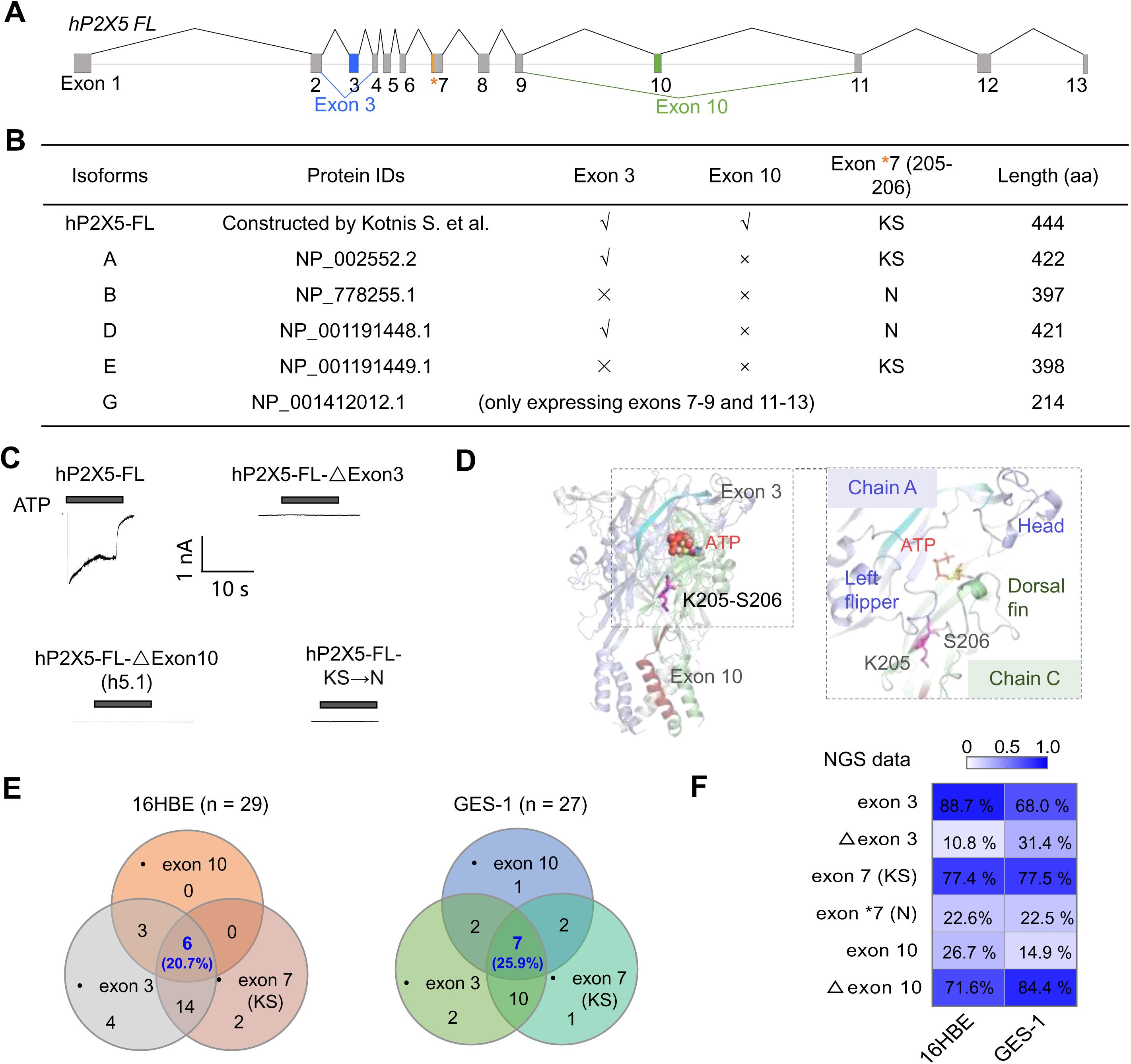
Gene expression and functional characterization of P2X5 in human cell lines. **(A)** Exon distribution patterns of different h*P2X5* transcripts. (**B)** Sequence characteristics of hP2X5 isoforms. (**C)** Representative current traces for hP2X5 WT and its mutants. (**D)** Homology model of hP2X5-ATP complex showing the positions of amino acids encoded by exons 3, 10, and the K205-S206 residues (encoded by exon 7). (**E)** Venn diagrams showing the number of hP2X5 T-A clones containing exons 3, 10, and full exon 7 (K205-S206) in 16HBE and GES-1 cell lines. Overlapping regions indicate clones containing two or three exons. (**F)** Heatmap displaying the inclusion ratios of exons 3 and 10, as well as full-length exon 7, in hP2X5 expressed in 16HBE and GES-1 cell lines. Data derived from NGS sequencing of hP2X5 PCR products.

To investigate the natural expression of exon 10-containing hP2X5, we cloned the *hP2RX5* gene from two human cell lines: normal human bronchial epithelial cells (16HBE) and gastric epithelial cells (GES-1). We observed high expression levels for both exon 3 and exon 7, with expression rates of 93.1% and 77.8% for exon 3, and 75.9% and 74.1% for exon 7 in 16HBE and GES-1, respectively (Fig. 3E and Table EV7). Exon 10 was expressed in approximately 30% of the clones (31.0% in 16HBE and 44.4% in GES-1) (Fig. 3E and Table EV6). Furthermore, around 20% of the clones co-expressed exons 3, 7, and 10 (20.7% in 16HBE and 25.9% in GES-1) (Fig. 3E), indicating that the full-length, ATP-responsive hP2X5-FL receptor is indeed expressed, albeit at lower levels than isoforms from other species such as zfP2X5, rP2X5, and mP2X5.

NGS was employed to further validate the expression of exons 3, 7, and 10. The NGS results confirmed that exon 3 and exon 7 were expressed at high levels, while the expression of exon 10 was reduced relative to monoclonal data (16HBE and GES-1 exon 3 expression rates were 88.7% and 68.0%, and exon 10 expression rates were 26.7% and 14.9%, respectively) (Fig. 3F). Despite the reduced expression of exon 10, it was still detectable in both cell lines. In addition, qPCR analysis confirmed the presence of exon 10 expression with copy numbers of 1.75×10³ ± 0.06 and 0.55×10³ ± 0.04/ng DNA for 16HBE and GES-1, respectively (n = 4, Fig. 3G). These results provide strong evidence that the full-length, ATP-responsive hP2X5 receptor is expressed in human cell lines. These findings challenge the previous assumption that such receptors exist solely in weak or non-responsive forms in species such as humans, rats, and mice (Collo et al., 1996, Cox et al., 2001, Le et al., 1997), and suggest that strong ATP responses may not be confined to a narrow subset of African populations (Kotnis et al., 2010). However, further studies are required to delineate the specific physiological, pathological, and developmental contexts in which these responses occur.

### Exploring the expression of exons 3, 7, and 10 of hP2X5 through RNA-Seq data from normal human tissues in the NCBI database

We analyzed RNA-Seq data from normal human tissues available in the NCBI database (PRJNA280600), which includes 20 tissue samples, to explore the expression of exons 3, 7, and 10 of hP2X5 (Fig. 4A-D). In the heart, no exon skipping event was observed in exon 3, which showed a 100% inclusion level (IncLevel = 1, Fig. 4A). In the spleen, which exhibited the highest *P2RX5* gene expression, exon 3 was largely retained (IncLevel = 0.96, Fig. 4A), with substantial retention in other tissues as well, including thymus, brain, trachea, fetal liver, stomach, and fetus (IncLevel values of 0.79, 0.74, 0.74, 0.70, and 0.64, respectively, Fig. 4A). Interestingly, although gene expression in fetal liver was low, exon 3 retention was still detected (IncLevel = 0.70 for fetal liver, Fig. 4A).

**Fig. 4.**
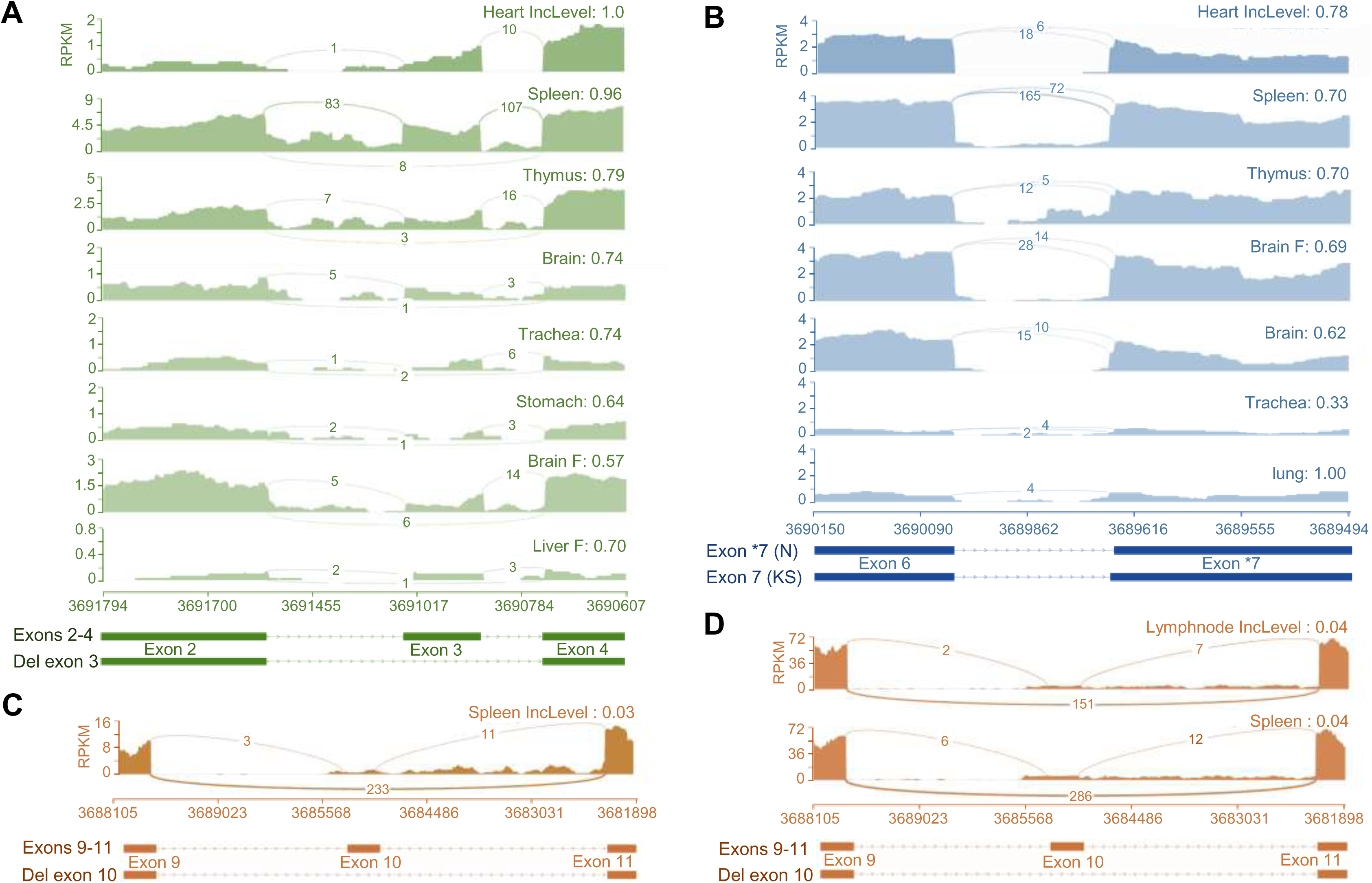
Analysis of alternative splicing of *P2X5* in RNA-Seq data from normal human tissues. Representative Sashimi plots showing RNA-seq read coverage of alternatively spliced exons 3 (A), 7 (B), and 10 (C, D) in human tissue samples. The density plot represents reads per kilobase per million mapped reads (RPKM) at each genomic position. Arcs represent splicing junctions, and numbers on the lines indicate junction-spanning reads. IncLevel indicates exon inclusion levels. The x-axis coordinates correspond to the gene location on the chromosome. Bottom sections show schematic diagrams of exon structures. Brain F and liver F represent fetal brain and fetal liver tissues, respectively.

In parallel, we identified alternative splicing events at the 3’ end of exon 7 (A3SS) in most tissues. In those with higher P2X5 expression, functional exon 7 (KS at residues 205-260) was predominantly retained, with inclusion levels exceeding 60% in tissues such as heart, spleen, thymus, fetal brain, and brain (IncLevel = 0.78, 0.70, 0.70, 0.69, and 0.62, respectively, Fig. 4B). In contrast, in the lung, where P2X5 expression was low, all transcripts retained functional exon 7, whereas in the trachea, only 33% of the transcripts contained functional exon 7 (IncLevel = 1.00, 0.33 for lung and trachea, respectively, Fig. 4B). While the lower expression in these tissues could lead to fewer splicing events, the data still provide compelling evidence of exon 7’s involvement in A3SS, with tissue-specific variations in inclusion levels.

Notably, exon 10 was found to undergo exon skipping in the spleen, with an inclusion level of 3% (Fig. 4C). In an additional dataset from 96 samples across 27 tissues (RRJEB4337), low levels of exon 10 retention were also observed in the spleen and lymph node (IncLevel = 0.04 for both tissues, Fig. 4D). Despite the low inclusion levels, the detection of exon 10, even at such reduced levels, emphasizes its possible role in regulating the function of the hP2X5 receptor in physiological conditions.

### Strong ATP Responsiveness of full-length P2X5 receptors across mammalian and non-mammalian species

In our comprehensive analysis, we assessed the ATP responsiveness of full-length P2X5 receptors from a range of species, including previously characterized organisms such as chicken and bullfrog [14, 16], as well as three unreported mammalian species — dog, bovine, and *Heterocephalus glaber* — using electrophysiological recordings in HEK293 cells. The data demonstrated that full-length P2X5 receptors from all species tested were capable of eliciting significant ATP-induced currents, with the following observed values: 13.9 ± 6.4, 120 ± 12, 107 ± 17, and 197 ± 7 pA/pF for cP2X5, fP2X5, dP2X5, bP2X5, and hgP2X5, respectively (n = 4-10; Fig. 5A, B). Of particular interest, the bovine P2X5 receptor showed ATP responsiveness only in its full-length, with a notable reduction in ATP current when exon 12 was absent (2.58 ± 0.60 pA/pF). In *Heterocephalus glaber* P2X5, a selective splicing event at the 5’ end of exon 4 resulted in the deletion of 11 amino acids, leading to a considerable loss of receptor functionality (17.4 ± 10.4 pA/pF; Fig. 5A, B).

**Fig. 5.**
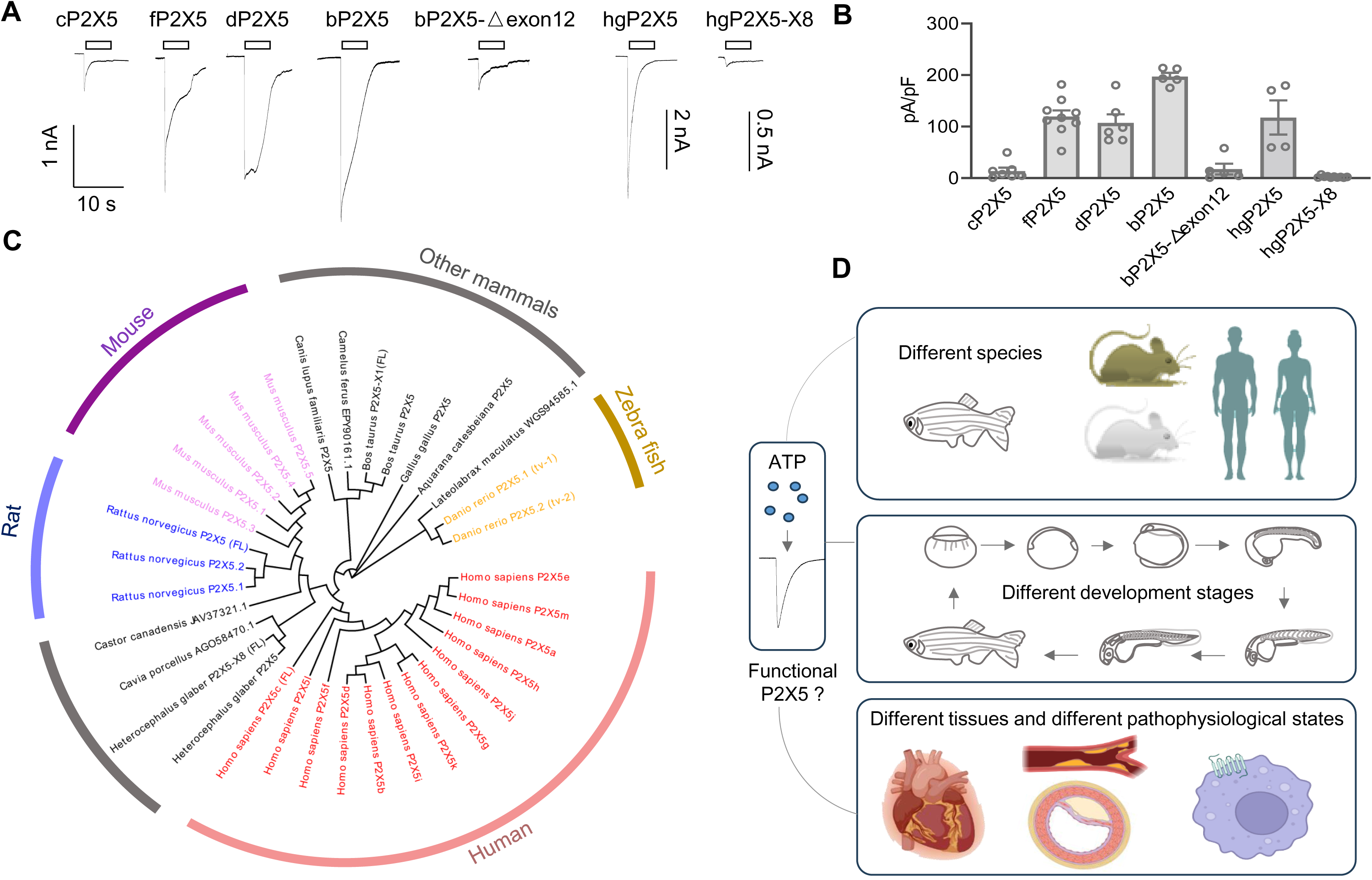
Phylogenetic analysis and functional validation of P2X5 expression across species. **(A-B)** Representative current traces (A) and pooled data (B) for cP2X5, fP2X5, dP2X5, bP2X5, hgP2X5, and their respective isoforms. Data are presented as mean ± SEM (n = 4-10). (**C)** Phylogenetic tree of P2X5 receptors, with corresponding accession numbers: zfP2X5.1 (tv-1), AAM74150.1; zfP2X5.2 (tv-2), AAI62598.1; rP2X5(FL), CAA65993.1; rP2X5.1, XP_006246628.1; rP2X5.2, XP_063124335.1; mP2X5.1, NP_201578.2; mP2X5.2, NP_001363911.1; mP2X5.3, NP_001363912.1; mP2X5.4, NP_001411762.1; mP2X5.5, NP_001412017.1; hP2X5a, NP_002552.2; hP2X5b, NP_778255.1; hP2X5c (FL), PQ800136; hP2X5d, NP_001191448.1; hP2X5e, NP_001191449.1; hP2X5f, PQ800137; hP2X5g, NP_001412011.1; hP2X5h, PQ800138; hP2X5i, PQ800139; hP2X5j, PQ800140; hP2X5k, PQ800141; hP2X5l, PQ800142; hP2X5m, PQ800143; hgP2X5, EHB05794.1; hgP2X5-X8 (FL), XP_004857285.1; bP2X5, NP_001291502.1; bP2X5-X1 (FL), XP_005220317.1; cP2X5, NP_990079; fP2X5, AAL24075.1; dP2X5, XP_038533703.1; Cavia porcellu P2X5, AGO58470.1; Castor canadensis P2X5, JAV37321.1; Camelus ferus P2X5, EPY90161.1; Lateolabrax maculatus P2X5, WGS94585.1. P2X5 receptors from human, mouse, rat, zebrafish, and other mammals are highlighted in light pink, purple, slate blue, olive, and gray, respectively. (**D)** A schematic representation illustrating the isoforms of the P2X5 receptor across various species, developmental stages, tissues, and pathophysiological states, and their responses to ATP, addressing the central question of this study.

Thus, our results show that full-length P2X5 receptors consistently mediate strong ATP responses across all species tested. In particular, high levels of expression of full-length P2X5 receptors were observed in various tissues and developmental stages in zebrafish, mouse, and rat, directly challenging the conventional belief that P2X5 receptors in mammals are predominantly weak or non-responsive to ATP (Bo et al., 2003, King, 2023, Kotnis et al., 2010, Le et al., 1997). To further investigate the molecular features underlying these observations, we conducted a phylogenetic analysis and sequence alignment of the verified functional isoforms of P2X5 (Fig. 5C), which includes sequence on the various isoforms of the P2X5 receptor that we obtained in this study. This analysis revealed the existence of novel splice variants and mutations that may contribute to the observed diversity in P2X5 receptor function, assisting in future studies into the functionality of P2X5 receptors. These isoforms likely exhibit differential expression in various tissues and at different developmental stages, and might undergo changes during disease progression, potentially contributing to specific functional outcomes. Given the potential pathophysiological significance of these findings, further investigation into the functional roles of these ATP-responsive P2X5 isoforms is essential (Fig. 5D).

## Discussion

Among the various P2X receptor subtypes, investigations into P2X5 receptors—both structural and functional—has lagged considerably, hindering a comprehensive grasp of their biological significance (Hattori & Gouaux, 2012, King, 2023, Saul et al., 2013, Schmid & Evans, 2019). Notably, P2X5 receptors are abundantly expressed in tissues such as the nervous, immune, muscular, and cardiovascular systems, highlighting their potential physiological and pathological importance (Coddou et al., 2011b, Illes et al., 2021, King, 2023). There is growing evidence that P2X5 may be involved in processes like inflammatory bone loss (Burnstock & Verkhratsky, 2010), immune regulation (Burnstock, 2008), cancer cell metastasis (Sun et al., 2019), and cellular proliferation and differentiation (Kim, Kajikawa et al., 2018), with implications for tumorigenesis and progression (Kim, Walsh et al., 2017). These diverse pathophysiological roles imply that P2X5 may harbor crucial yet insufficiently explored functional characteristics, underscoring the need for deeper investigation into their molecular mechanisms. Here, we examined the ATP response mechanisms of various splice isoforms and transcriptional variants of P2X5 receptors from several species. Our initial focus was on two zebrafish P2X5 transcriptional variants, zfP2X5^TV1^ and zfP2X5^TV2^, which exhibit substantial functional differences(Kucenas et al., 2003, Low et al., 2008). We employed saturation mutagenesis, receptor membrane expression assays, conductance measurements, along with kinetic and conformational sampling of mutant forms to identify the key residues that modulate the functional properties of zfP2X5. Our results indicate that these critical sites are located in non-conserved regions that play pivotal roles in P2X gating and ATP recognition, suggesting that the expression of zebrafish P2X5 may regulate the intensity of ATP responses through mutations in these hypervariable regions. Intriguingly, functional analysis revealed that the zfP2X5^TV2^ isoform, despite carrying high-frequency mutations (K213N and an insertion mutation 397-401), consistently exhibited strong ATP responsiveness. Furthermore, RNA sequencing analyses of developmental stages and high-expression tissues from rats and mice, revealed that the rat and mouse primarily express ATP-responsive mP2X5^G317^ (> 90%) and rP2X5^F191^ (> 70%), respectively. Despite a reduced frequency, sequencing of human airway epithelial cell line 16HBE and gastric epithelial cell line GES-1 mRNA by T-A cloning and NGS sequencing detected full-length exons 3, 7, and 10 of hP2X5 in 15-30% of cases, underscoring their essential role in maintaining strong ATP responsiveness. RNA-seq analysis of human tissue samples from the NCBI database (RRJEB4337, including 96 samples from 27 tissues) revealed a higher expression of exons 3 and 7, whereas exon 10 was less prevalent in certain local organs, suggesting polymorphisms or the existence of multiple transcriptional variants of hP2X5. The potential underestimation of this data might be attributed to insufficient deep sequencing of normal human samples in current databases. Additionally, we assessed the ATP responses of full-length P2X5 sequences from species such as chicken, bullfrog, dog, cow, and naked mole rat, all of which exhibited strong ATP-induced currents. These results prompted us to reconsider the prevailing view that “P2X5 homotrimers are typically weak or non-responsive to ATP “(Bo et al., 2003, Collo et al., 1996, King, 2023, Kotnis et al., 2010, Le et al., 1997). Indeed, P2X5 receptors are highly expressed in tissues such as the nervous, immune, skeletal, and cardiovascular systems across most mammals, yet exhibit significant variation in ATP responsiveness among their splice variants and transcriptional forms. These findings emphasize the importance of reconsidering P2X5 isoforms that exhibit strong ATP responsiveness.

In our study, we analyzed P2X5 receptors from zebrafish, mice, and rats, uncovering more frequent detection of ATP-responsive isoforms and mutants across a range of developmental stages and tissues. For example, during zebrafish development, the strong ATP-responsive zfP2X5^TV2^ isoform was predominantly expressed, while the ATP-insensitive zfP2X5^TV1^ isoform, identified in previous studies (Diaz-Hernandez, Cox et al., 2002), was not detected. Similarly, in both mice and rats, ATP-responsive P2X5 forms were consistently detected throughout their lifespan. These apparent discrepancies can likely be attributed to the methodological limitations of earlier studies, or perhaps the transient nature of non-ATP-responsive isoforms during specific developmental windows or in particular tissues. Early cloning studies of P2X5 receptors from rats, mice, and humans in the 1990s (Collo et al., 1996, Cox et al., 2001, Le et al., 1997) were constrained by the technological limitations of the time, which hindered both the accuracy and depth of sequencing, resulting in an incomplete cataloging of P2X5 isoforms. Consequently, the receptor was erroneously considered non-functional, leading to limited subsequent investigation (Illes et al., 2021, King, 2023). However, with recent advancements in sequencing and transcriptomic technologies (Stark et al., 2019, Trapnell, 2024), fresh insights have emerged, although updated sequences of P2X5 have not garnered the attention they deserve, potentially resulting in the oversight or misinterpretation of key functional isoforms. Furthermore, while earlier studies suggest that ATP-insensitive forms of P2X5 may exist under specific conditions (Bo et al., 2003, King, 2023, Kotnis et al., 2010, Le et al., 1997), their distribution in time and space is likely highly restricted. Studies have indicated that gene splicing or mutations may transiently occur during certain developmental stages or within specific tissues (Bo et al., 2003, de Rijke, van Horssen-Zoetbrood et al., 2005, Duckwitz, Hausmann et al., 2006, King, 2023, Kotnis et al., 2010, Lê, Boué-Grabot et al., 1999, Overes, De Rijke et al., 2008, Schiller et al., 2022), leading to a temporary loss of receptor functionality. While such selective exon splicing or mutations may occur sporadically, their frequency remains low (King, 2023, Kotnis et al., 2010, Overes, Fredrix et al., 2009). As a result, these forms are likely of limited physiological relevance and unlikely to represent the primary functional state of the receptor. Our studies, focusing on the predominant forms, captures the biologically significant P2X5 isoforms that play crucial roles across various tissues and developmental stages. While ATP-insensitive forms may exist, their rarity and transient nature likely make them difficult to detect experimentally.

Furthermore, our analysis reveals that the full-length, strongly ATP-responsive form of human P2X5 was less frequently detected in normal human cell lines and tissue samples than in non-human mammals and non-mammalian species. This finding may reflect an evolutionary adaptation of hP2X5 to meet the specialized needs of higher organisms, particularly within the immune and nervous systems (Donnelly-Roberts, Namovic et al., 2009, King, 2023, Serrano, Mo et al., 2012, Sun et al., 2019). The low frequency of strong ATP-responsive hP2X5 forms in humans could indicate a greater degree of functional diversification, suggesting a more complex regulatory role for the receptor in human physiology compared to that in lower organisms. In contrast to non-mammalian species and non-human mammals, hP2X5 may govern more intricate biological processes such as immune modulation and neuroendocrine regulation. P2X family receptors are integral to inflammation, cellular signaling, and neurotransmitter release—processes that vary substantially between humans and rodents. As such, hP2X5 likely operates within a more elaborate regulatory framework, where splice variants or mutations fine-tune immune responses and central nervous system functions (Hattori & Gouaux, 2012, Illes et al., 2021, King, 2023, Lalo et al., 2008). It is also important to consider that the detection frequency of hP2X5 may be influenced by the experimental models and data analysis methods employed. The normal cell lines studied, including 16HBE and GES-1, likely represent limited physiological states, with the observed 20% detection rate of strong ATP-responsive P2X5 forms potentially reflecting a cell line-specific occurrence rather than an accurate representation of systemic expression. Moreover, RNA-seq data from public databases, which are often derived from samples collected across different contexts and time points, may contain discrepancies that affect the precise quantification of P2X5 expression across tissues and developmental stages.

Thus, our findings indicate that strong ATP-responsive P2X5 isoforms may play a role in several biological processes, with their functional properties likely influencing the physiological activities of diverse systems, including the nervous and immune systems (Coddou et al., 2011b, Illes et al., 2021, King, 2023). This new understanding challenges the traditional view of P2X5 as a non-functional receptor, urging a reevaluation of the existing findings and database descriptions of P2X5. To advance our understanding, further investigations employing cutting-edge technologies, such as single-cell transcriptomics (Wu, Yang et al., 2024) and spatial genomics (Gulati, D’Silva et al., 2025), will offer more precise insights into the temporal and spatial distribution of P2X5 isoforms. Additionally, a deeper exploration of how selective splicing events and mutations modulate P2X5 function is crucial for uncovering its role in various pathophysiological processes. These efforts will not only enrich our understanding of P2X5 but also enhance our ability to elucidate its contribution to both normal physiology and disease.

## Materials and methods

### Plasmids and mutagenesis

Plasmids for rP2X5 and hP2X5 were provided by Drs. Lin-Hua Jiang and Alan North. cDNAs for mP2X5 isoforms (5.1, 5.2, 5.3) were cloned from C57BL/6 mice and inserted into pCDNA3.1. cDNAs for zfP2X5-TV2, cP2X5, fP2X5, dP2X5, bP2X5-FL, hgP2X5-FL, and hgP2X5-X8 were synthesized by BGI and subcloned into pEGFP-N3. Other plasmids were generated using NEBuilder® HiFi DNA Assembly (NEB) or KOD-Plus Mutagenesis Kit (Toyobo).

### Cell and molecular biology

Unless otherwise specified, all compounds were obtained from Sigma-Aldrich (USA). The 16HBE14o-human bronchial epithelial cells (16HBE) and human gastric epithelial cells (GES-1) were generously provided by Dr. Ying-Zhe Fan. Human embryonic kidney (HEK293) cells (Catalog ID: GNHu 43) were purchased from the Shanghai Institutes for Life Sciences, Chinese Academy of Sciences, Cell Repository. The cells were cultured in Dulbecco’s Modified Eagle Medium (DMEM) supplemented with 10% fetal bovine serum, 1% penicillin/streptomycin, and 1% GlutaMAX, and were maintained at 37°C in a humidified atmosphere with 5% COL. Wild-type (WT) and mutant plasmids were transfected into HEK293 cells using calcium phosphate precipitation, as previously described (Wang, Zhang et al., 2024).

### Electrophysiological recordings

Electrophysiological recordings were performed 24 to 48 hours post-transfection at room temperature (25 ± 2°C) using an Axopatch 200B amplifier (Molecular Devices) in voltage-clamp mode, with the holding potential set to −60 mV (Yang, Niu et al., 2017). Patch pipettes were fabricated from 1.5 mm borosilicate glass capillaries using a two-stage puller (PC-10, Narishige, Japan), yielding a resistance range of 3-5 MΩ. Current signals were acquired in Gap-free mode with a sampling rate of 10 kHz and filtered at 2 kHz, with data analyzed using pClamp 10.0 (Molecular Devices). The extracellular solution contained the following (in mM): 150 NaCl, 5 KCl, 10 glucose, 10 HEPES, 2 CaClL·2HLO, and 1 MgClL·6HLO, with the pH adjusted to 7.35 –7.40 using Tris-base. Patch pipettes were filled with the following internal solutions (mM): for whole-cell recordings, 120 KCl, 30 NaCl, 1 MgClL·6HLO, 0.5 CaClL·2HLO, 10 HEPES, and 5 EGTA (pH 7.35–7.40); for nystatin-perforated patch-clamp recordings, 75 KLSOL, 55 KCl, 5 MgSOL, and 10 HEPES. Nystatin-perforated patch-clamp recordings (0.15 mg/mL nystatin) were employed to prevent current rundown during repetitive ATP applications, with ATP applied at approximately 8-minute intervals. Inhibitors were pre-applied for 1 minute before co-application with ATP. For single-channel recordings, recording electrodes were pulled from borosilicate glass (World Precision Instruments, Inc.), with a resistance of approximately 10 MΩ. To facilitate larger single-channel currents, the holding potential was set to −120 mV. Data were sampled at 50 kHz and filtered at 2 kHz. The external solution for single-channel recordings was free of calcium and magnesium, matching the conditions used for whole-cell recordings.

### Homology modeling and molecular simulations

Homology models for zfP2X5^TV2^, rP2X5, mP2X5 and hP2X5 were constructed using the crystal structures of zfP2X4 in both the *apo* and open states (PDB IDs: 4DW0 and 4DW1) as templates. Sequence alignments were performed using ClustalW, and homology modeling was carried out with Modeler 9.9 (Chovancova, Pavelka et al., 2012). The top-scoring models were validated using ProCheck (Roman, Laskowaski et al., 1993). To model the membrane environment, a POPC bilayer was incorporated into the transmembrane region of P2X5 using the System Builder module in DESMOND (Dror, Arlow et al., 2011). The system was solvated in SPC (simple point charge) water with 150 mM NaCl, and counterions were added to neutralize the charge. Following default relaxation with the NPT ensemble (constant pressure, constant temperature, and constant particle number), a 0.5-μs molecular dynamics (MD) simulation was run, with trajectory recording intervals set at 200 ps. The all-atom OPLS_2005 force field was applied (Banks, Beard et al., 2005, George, Kaminski et al., 2001, William, Jorgensen et al., 1996). Simulations were carried out on a DELL T7920 workstation equipped with an NVIDIA Tesla K40C or CAOWEI 4028GR with an NVIDIA Tesla K80 GPU.

### Western Blotting

HEK293 cells were transfected with either wild-type or mutant P2X5 plasmids, washed three times with phosphate-buffered saline (PBS, pH 7.4), and incubated with 2 mL of sulfo-NHS-LC-biotin (Pierce, Germany). The cells were placed on ice at 4°C for 30 minutes, with intermittent shaking every 10 minutes to ensure uniform labeling of membrane-bound proteins. Glycine was added to terminate the reaction. Following three additional washes with PBS, the cells were lysed in 200 μL of RIPA buffers. The lysate was collected after scraping the cells from the culture dish and centrifuged at 12,000 rpm for 30 minutes at 4°C. A 20% (v/v) aliquot of the supernatant was mixed with SDS loading buffer for total protein quantification. The remaining supernatant was incubated overnight at 4°C with NeutrAvidin-conjugated agarose resin (Thermo Scientific, USA), washed five times with cold PBS, and eluted in SDS loading buffer. Protein samples were separated by SDS-PAGE in a metal bath for 10 minutes before being transferred to polyvinylidene difluoride (PVDF) membranes (Immobilon-P). The membranes were blocked with 5% skim milk (Difco) for 2 hours at room temperature and then incubated overnight at 4°C with primary antibodies (anti-FLAG, 1:3000; Sigma-Aldrich or anti-GAPDH, 1:3000; Sungene Biotech, China), both in 5% skim milk. After washing, membranes were incubated with horseradish peroxidase-conjugated secondary antibodies (goat anti-mouse IgG(H+L)-HRP, 1:3000; Sungene Biotech, China) at 25 ° C for 2 hours. The protein expression was visualized using electrochemiluminescence (ECL) reagents (HIGH, Thermo Scientific, USA) on the ImageQuant RT-ECL system (Tanon 5200, China). The analysis was repeated in at least three independent experiments.

### *P2RX5* gene amplification, sequencing and analysis Animals

C57BL/6 mice (catalog ID: SM-001) and SD rats (catalog ID: SCSP-402) were purchased from Jiangsu Huachuang Sino (China). All animals were housed under controlled conditions (23 ± 2°C, 50 ± 10% humidity, 12-hour light/dark cycle) with ad libitum access to standard food and water. Zebrafish (Sanger AB Tübingen line, catalog ID: CZ1) were a generous gift from Dr. Jing Xiang and maintained in freshwater (pH 7.0-8.0, 28 ± 2°C, 14-hour light/10-hour dark cycle) with standard food.

### Tissue preparations

For zebrafish, embryos were collected at 0-hour post-fertilization (hpf), and then at 6, 12, 24, 36, and 48 hpf. Adult zebrafish (10-12 weeks old) were used for tissue collection. In mice and rats, the presence of a vaginal plug was designated as embryonic day 0. Tissues (heart and brain) were collected at embryonic day 17 (E17) for mice and E18 for rats, as well as at postnatal day 0 (P0), P21, and in adult animals (6-8 weeks old). All animals were euthanized by asphyxiation using a rising concentration of CO_2_.

### Gene amplification

RNA was extracted using TRIzol and reverse transcribed into cDNA with the PrimeScript RT Reagent Kit (TaKaRa). The P2RX5 gene was amplified with PrimeSTAR HS DNA Polymerase (TaKaRa), and PCR products (1,000-2,000 bp) were purified and recovered. The following primer sets were used for amplification:

*zfP2RX5* forward: CTTCCAGGCTGTCATTGA, reverse: ACTGGCATTATTCCTCATTG; *mP2RX5* forward: AAGTGAGCTGGAGGCATGGG, reverse: GTTCCCTGCCAAGAGCTCT; *rP2RX5* forward: CGGCAGCGCACACAGAGT, reverse: GGCTACGTCTTCACTGGATGC; *hP2RX5* forward: CTGAGAGCGCGCCATGG, reverse: CCTGAACGTAAGCAGAGGC.

The target sequences were amplified, and PCR products were purified for T-A cloning and next generation sequencing (NGS) analysis.

### T-A cloning

Purified PCR products (1,000-2,000 bp) were treated with an “A” tail at the 3’ end and cloned into a T-vector (TaKaRa). Primers for sequencing and subsequent analysis were synthesized by GENEWIZ.

### NGS analysis

Fragments of 1500-2000 bp are purified and recovered. All sequencing samples were prepared and submitted to Genewiz for sequencing services. Upon receipt, the company conducted quality control and purification of the samples, followed by next generation amplicon library construction and subsequent high-throughput sequencing on the Novaseq 2×150 bp platform. Subsequently, the raw data underwent statistical analysis, with Cutadapt (1.9.1) used to remove adapters and low-quality sequences (Martin, 2011). The quality-controlled data were merged using Pandase (2.7) for clean reads (Masella, Bartram et al., 2012). After data demultiplexing based on barcodes, the sequence counters (SeqCount) and base distribution (BaseDist) of the target sequences were analyzed.

### Real-time quantitative PCR (qPCR)

qPCR was conducted using the standard curve method for absolute quantification. A 10-fold serial dilution of plasmid containing the *P2RX5* gene was used to generate a standard curve for real-time qPCR amplification using TB Green Premix Ex Taq (TaKaRa). The following primers were used to analyze the expression of *P2RX5* genes:

*zfP2RX5* forward: TGTCCTGCCTATCACCAA, reverse: CCACTCAATTCCAATTCCAA; *mP2RX5* forward: GCTCCACCAATCTCTACTG, reverse: GTTGTCCAGACGGTTGAA; *rP2RX5* forward: CGAGTGTTCTGAGGATGAT, reverse: GGAAGCGAATGAAGTTCTT; *hP2RX5* forward: CTGGTCGTATGGGTGTTCCT, reverse: CTGGGCTGGAATGACGTAGT.

For exon 10 expression in human *P2RX5*, the forward primer was TTGCCAGATATTACCGAGAC, and the reverse primer was GATGGTGGGAATGATGCT.

### Alternative splicing analysis

BioProject data for human (Accession: PRJNA267840, RRJEB4337) and rat (PRJNA238328) were downloaded from the NCBI database. Raw data quality was assessed with Trim Galore (v0.6.10) and aligned using STAR (v2.7.11b). Reference genomes for human (GRCh38.p14) and rat (GRCr8) were used. Alternative splicing events were analyzed using rMATS (v4.3.0), and relevant splicing diagrams were generated using rmats2sashimiplot(Wang, Xie et al., 2024).

### Data analysis

Data are presented as means ± SEM. Statistical analysis was performed using ANOVA or Student’ s t-test, with a p-value of < 0.05 considered statistically significant. Concentration-response relationships for ATP activation of WT or mutant channels were established by measuring currents in response to varying ATP concentrations. Data were fitted to the Hill equation: I/I_max_ = 1/[1 + (EC_50_/[ATP])^n^], where I represents the normalized current at a given concentration, I_max_ is the maximum normalized current, EC_50_ is the concentration eliciting half-maximal current, and n is the Hill coefficient.

## Acknowledgements

This work was supported with funds from Innovation and Entrepreneurship (Shuangchuang) Program of Jiangsu Province (JSSCTD202350), National Natural Science Foundation of China (No. 32371289), Natural Science Foundation of Jiangsu Province (BK20202002), Changsha “Jie Bang Gua Shuai” Major Science and Technology Programs (KQ2301004), “Xing Yao” Leading Scholars of China Pharmaceutical University (2021), the CAMS Innovation Fund for Medical Sciences (CIFMS) (2019-I2M-5-074) and the Medical Innovation and Development Project of Lanzhou University (lzuyxcx-2022-156).

## Author contributions

Y. Y. and X.N. Y. designed the project; C.H. S. and D.P. W. performed cell patch-clamp; C.H. S., D.P. W., H. Y. and X. C. performed cell culture, tissue preparations, gene amplification, T-A cloning, qPCR and Western blotting; H. Y., Y. Y. and X.N. Y. performed RNA-Seq data analysis; Y. Y. and X.N. Y. did homology models and MD simulations; Y. Y., X.N. Y., D.P. W. and H. Y., analyzed data; Y. Y., X.N. Y., D.P. W. and H. Y. wrote the manuscript. All authors discussed the results and commented on the manuscript.

Source data underlying figure panels in this paper may have individual authorship assigned. Where available, figure panel/source data authorship is listed in the NCBI and Uniport database.

## Data and material availability

No data have been deposited to public repositories.

The source data of this paper are collected in the NCBI and Uniport database.

## Expanded View Figure legends

**Fig. EV1.**
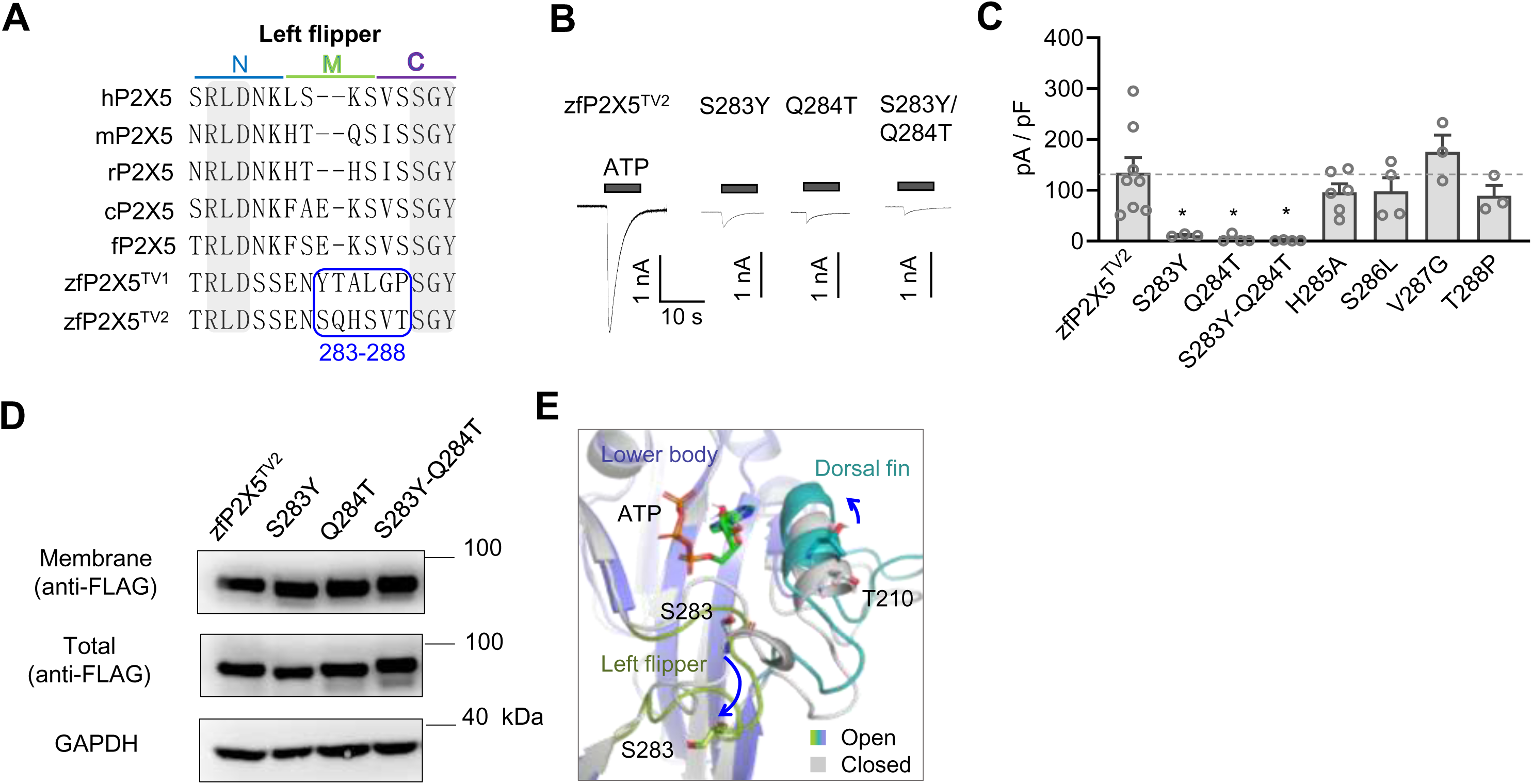
Functional significance of S283 and Q284 in the non-conserved left flipper domain of zfP2X5. **(A)** Sequence alignment of the loop 283-288 region in the left flipper domain between TV1 and TV2 of *zfP2X5*. (**B, C)** Representative current traces (B) and pooled data (C) for WT zfP2X5 and its mutants (S283Y, Q284T, S283Y/Q284T). Data are expressed as mean ± SEM (n = 3-8). Statistical significance: \**P* < 0.05 versus WT, one-way ANOVA followed by Dunnett’s multiple comparisons test, F(7, 27) = 5.76. (**D)** Representative Western blot images of WT zfP2X5-TV2 and its mutants. (**E)** Superimposition diagram illustrating the positional changes of S283 and T210 in the zfP2X5 receptor between its open and closed states. Blue arrows indicate positional shifts upon ATP binding.

**Fig. EV2.**
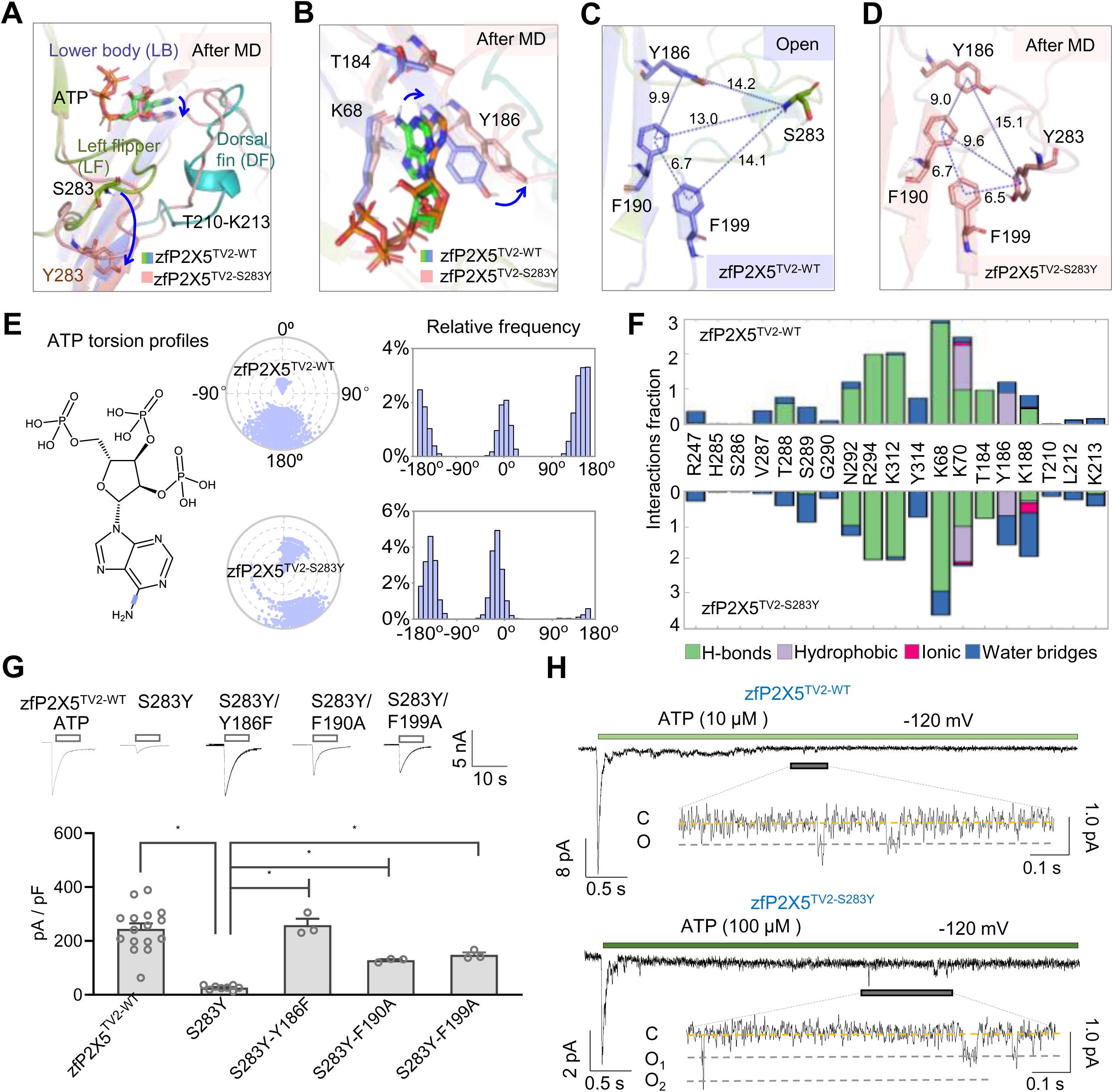
Residue S283 regulates ATP recognition in zfP2X5. (**A)** Downward motion of the dorsal fin and left flipper during the molecular dynamics (MD) simulation of zfP2X5-TV2 S283Y. **(B)** The Y186 residue near the binding pocket undergoes significant shifts and rotations following the mutation of residue 283 to Tyr. (**C, D)** Diagrams depicting the positional relationships between Y186, F190, F199, and either S283 (C) or Y283 (D). The introduction of Tyr at position 283 brings the other residues closer together. (**E)** In the chemical structure of ATP, key rotatable bonds are marked in purple. The radial plot shows conformational changes of the torsion bodies, with the histogram condensing the data representing the probability density of torsional angles. (**F)** Interaction dynamics between zfP2X5 and ATP, monitored throughout the simulations of both WT zfP2X5^TV2-WT^ and the zfP2X5^TV2-S283Y^ mutant. (**G)** Representative current traces (upper panel) and pooled data (lower panel) for zfP2X5 and its mutants. Data are expressed as mean ± SEM (n = 3-16). Statistical significance: *P < 0.05 versus WT, one-way ANOVA followed by Dunnett’s multiple comparisons test, F(4, 28) = 19.1; *P < 0.05 versus zfP2X5^TV2-S283Y^, one-way ANOVA followed by Dunnett’s multiple comparisons test, F(3, 13) = 124.1. (**H)** Representative current traces recorded from outside-out patches of wild-type zfP2X5^TV2-WT^ and the zfP2X5^TV2-S283Y^ mutant at a holding potential of −120 mV, in response to ATP concentrations of 10 μM and 100 μ M. All experiments were repeated at least three times.

**Fig. EV3.**
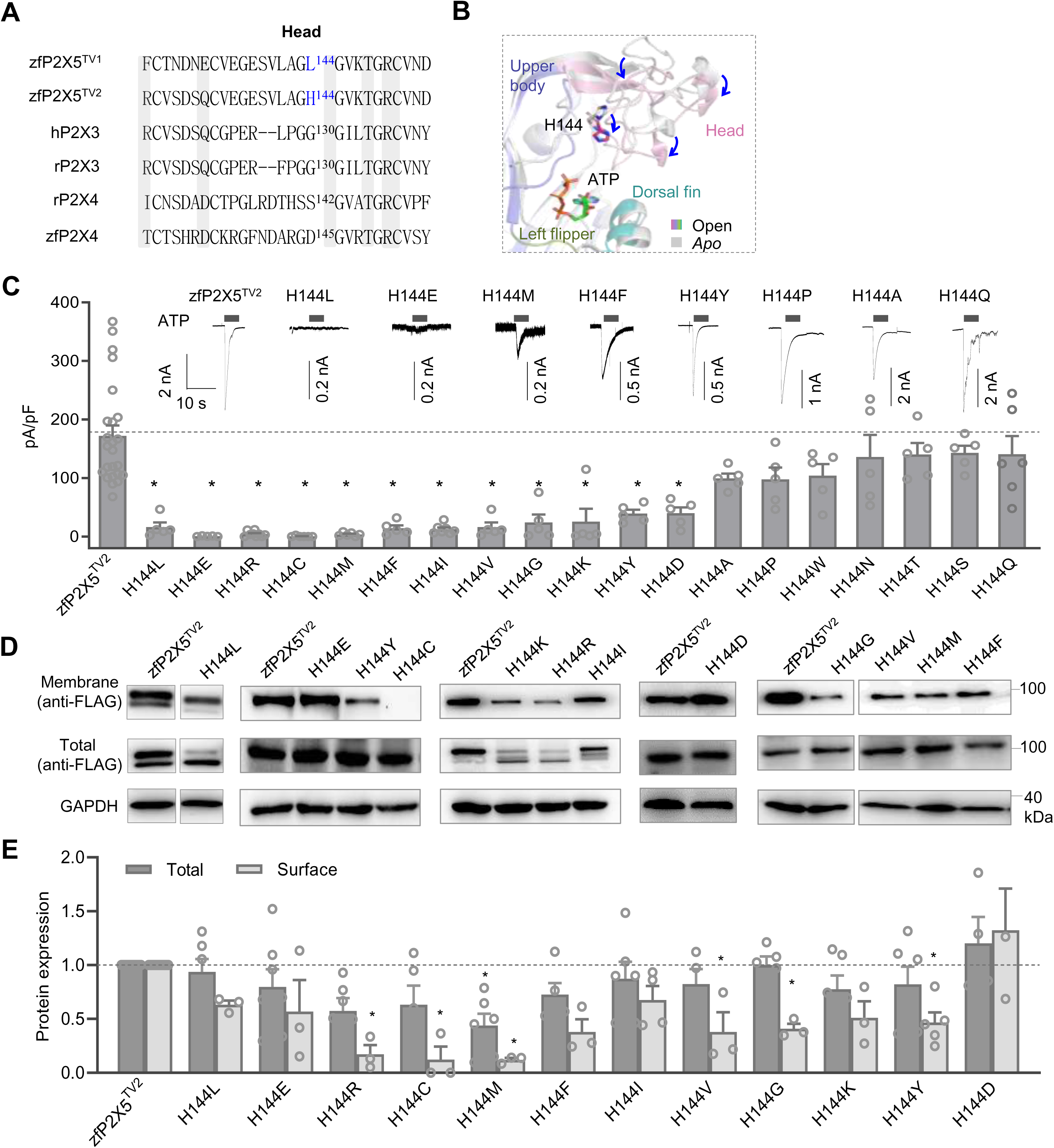
Substitution of residue H144 in the head domain alters the ATP response of the zfP2X5 receptors. (**A)** Sequence alignment of residue H144 in zfP2X5 isoforms (zfP2X5^-TV1^, zfP2X5^-TV2^) and homologous receptors in other species (*hP2X3*, *rP2X3*, *rP2X4*, and *zfP2X4*). (**B**) Superimposition diagram showing the positional changes of the head domain and residue H144 in the zfP2X5 receptor between the open and closed states. The blue arrows indicate downward movement upon ATP binding. (**C)** Representative current traces (upper panel) and pooled data (lower panel) for WT zfP2X5 and its mutants. Data are presented as mean ± SEM (n = 3-4). Statistical significance: *P < 0.05 versus wild-type, one-way ANOVA followed by Dunnett’s multiple comparisons test, F(19, 108) = 11.8. (**D-E**) Representative Western blot images (D) and pooled data (E) showing total and surface protein expression levels of zfP2X5 and its mutants. Data are presented as mean ± SEM (n = 3-15). Statistical significance: *P < 0.05 versus wild-type, one-way ANOVA with Dunnett’s multiple comparisons test, F(12, 74) = 2.90 and F(12, 40) = 8.81.

**Fig. EV4.**
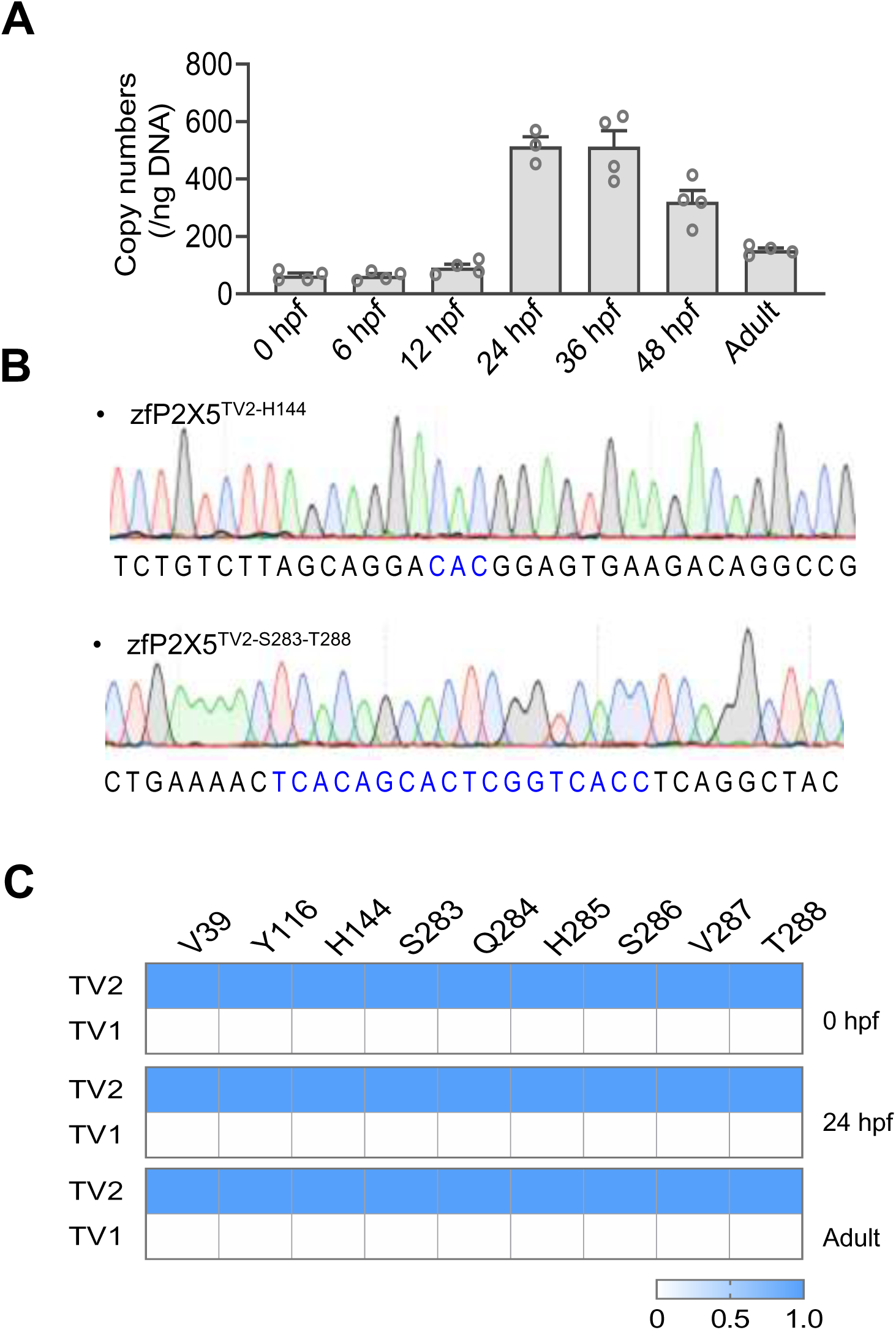
Expression analysis of the WT *zfP2RX5* gene sequence. (**A)** Quantitative PCR (qPCR) analysis of *P2RX5* gene expression at various developmental stages in zebrafish. (**B)** Sequencing diagram of the *P2X5* T-A clone. (**C)** Mutation analysis of the *zfP2X5* cloning product using next generation sequencing (NGS) data.

**Fig. EV5.**
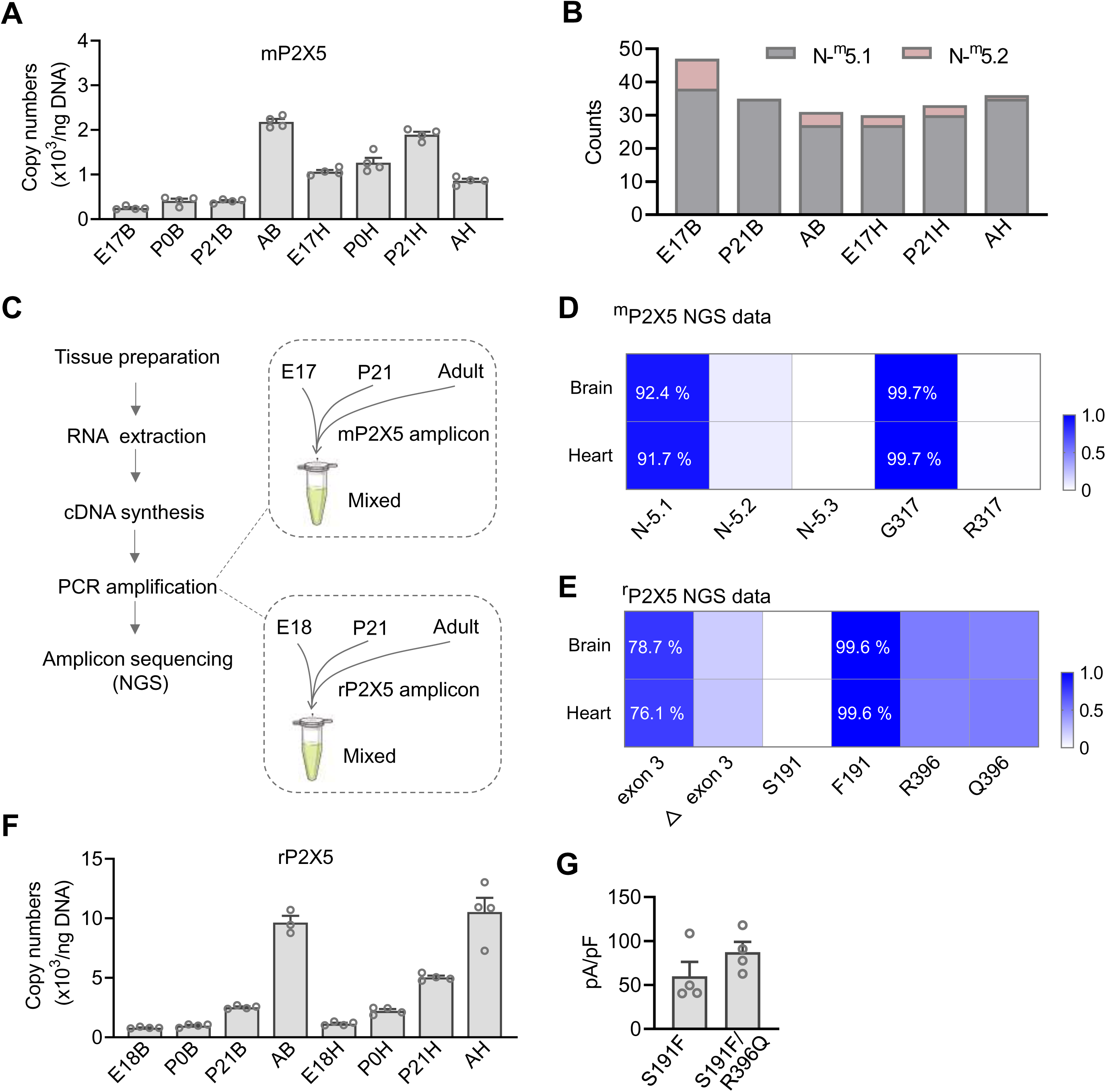
Expression analysis of the m*P2X5* and *rP2RX5* gene sequences. **(A)** qPCR analysis of *P2RX5* gene expression at various developmental stages in mice. (**B)** Statistical analysis of differential expression of *P2X5* splice variants across different developmental stages in the mouse. **(C)** Protocol for NGS-based genotyping and expression analysis of the *P2RX5* gene in mouse and rat. (**D-E**) Statistical analysis of the frequency of *P2X5* isoforms and mutations in various tissues of mice (D) and rat (E) across developmental stages based on NGS data. (**F)** qPCR analysis of *P2RX5* gene expression at various developmental stages in rat. (**G)** Aggregated current density data for *rP2X5* mutants S191F and S191F/R396Q. Statistical comparison: P > 0.05, unpaired t-test, S191F versus S191F/R396Q. Data are expressed as mean ± SEM (n = 3-4).

## Expanded View Tables

**Table EV1.**
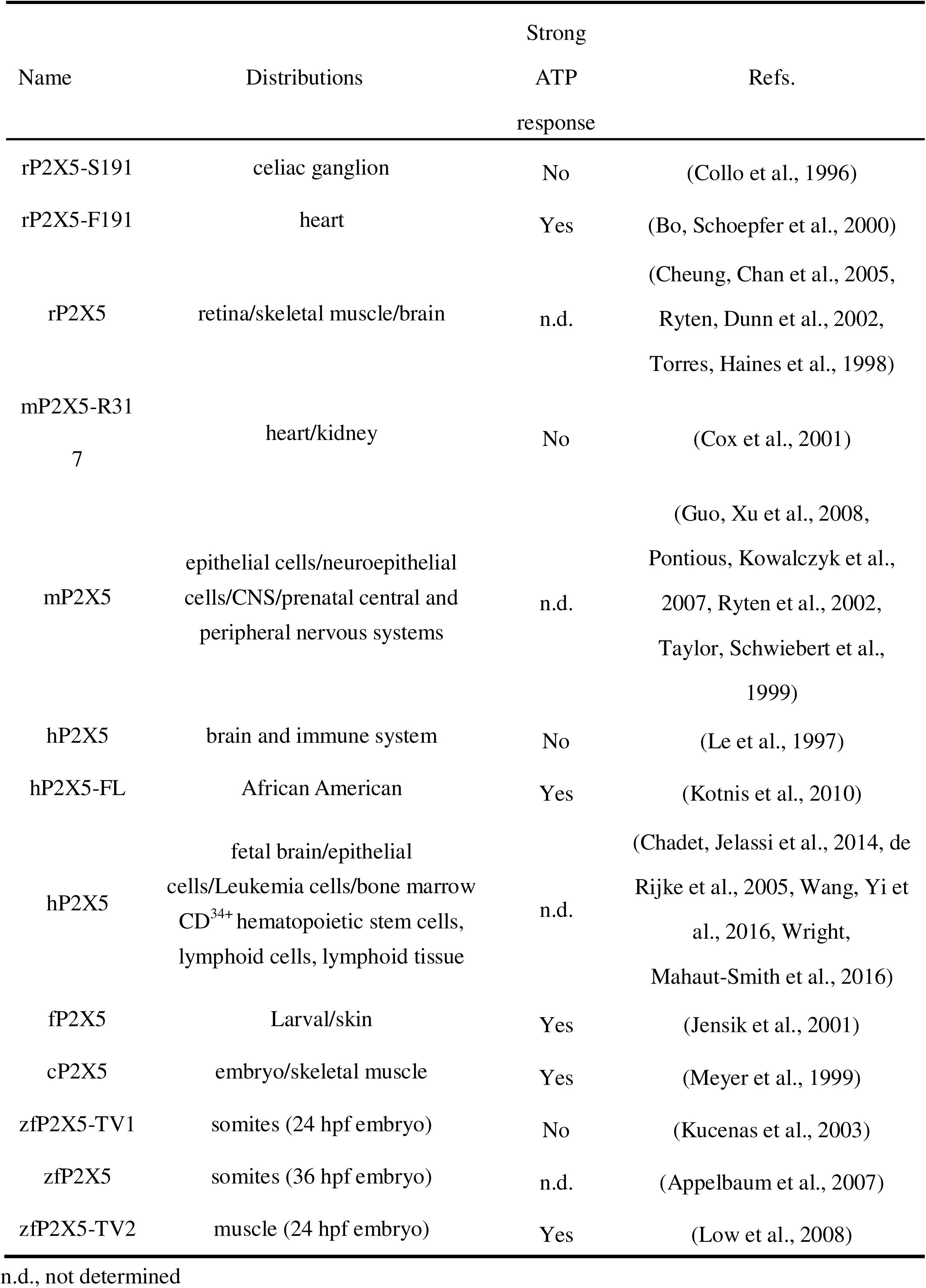
Expression and function of P2X5 receptors.

**Table EV2.** zfP2X5 mutations from adult, 0 hpf and 24 hpf.

| Mutation | Adult | 0 hpf | 24hpf |
| --- | --- | --- | --- |
| G96R | 0.46% | 0.02% | 0.01% |
| F127Vf | 0.04% | 0.90% | 0.05% |
| E133K | 0.40% | 0.03% | 0.04% |
| T157A | 0.39% | 0.01% | 0.02% |
| K179M | 0.59% | 0.11% | 0.13% |
| L204P | 0.44% | 0.02% | 0.02% |
| H267P | 0.01% | 0.01% | 0.54% |
| D278E | 0.33% | 0.30% | 0.38% |
| R294K | 1.11% | 1.68% | 0.31% |
| L344S | 0.35% | 0.01% | 0.02% |
| A351V | 0.25% | 0.42% | 0.27% |
| R396H | 2.70% | 2.61% | 2.85% |
| E426P | 0.43% | 0.02% | 0.14% |
| E464G | 0.01% | 0.44% | 0.01% |

**Table EV3.** Cloning of *mP2RX5* at different development stages.

| Period | Residues<br>317 | N-terminal |  |  |
| --- | --- | --- | --- | --- |
|  |  | N-5.1 | N-5.2 | N-5.3 |
| E17B | G (47/47) | 38 | 9 | n.d. |
| P21B | G (35/35) | 35 | 0 | n.d. |
| AB | G (31/31) | 27 | 4 | n.d. |
| E17H | G (27/27) | 27 | 3 | n.d. |
| P21H | G (33/33) | 30 | 3 | n.d. |
| AH | G (36/36) | 35 | 1 | n.d. |

n.d., not determined

**Table EV4.** Cloning of *mP2RX5* at different development stages.

n.d., not determined **Table EV4. Cloning of *mP2RX5* at different development stages**
| Periods | n | Indels (n) |
| --- | --- | --- |
| E17B | 47 | 2 |
| P21B | 35 | 10 |
| AB | 31 | 10 |
| E17H | 30 | 9 |
| P21H | 30 | 5 |
| AH | 35 | 6 |

**Table EV5.** Cloning of *rP2RX5* at different development stages.

| Period | n | 396R | R396Q | Indel |
| --- | --- | --- | --- | --- |
| E18B | 30 | 16 | 14 | 0 |
| P21B | 31 | 23 | 8 | 0 |
| AB | 30 | 16 | 14 | 0 |
| E18H | 30 | 5 | 16 | 9 |
| P21H | 36 | 16 | 19 | 1 |
| AH | 30 | 17 | 13 | 0 |

**Table EV6.** Cloning of *rP2RX5* at different development stages.

| Period | N | FL | Δexon3 | Δexon2/3 | Δexon 2/3/4 | Δexon 6 | Others |
| --- | --- | --- | --- | --- | --- | --- | --- |
| E18B | 30 | 14 | 1 | 12 | 1 | 0 | 5 |
| P21B | 31 | 17 | 1 | 8 | 4 | 3 | 3 |
| AB | 30 | 16 | 1 | 9 | 2 | 1 | 3 |
| E18H | 30 | 13 | 0 | 9 | 2 | 0 | 9 |
| P21H | 36 | 10 | 3 | 19 | 1 | 3 | 4 |
| AH | 30 | 21 | 2 | 6 | 0 | 0 | 1 |

**Table EV7.** The number of *hP2RX5* T-A clones that contain exons 3 and 10, as well as a complete exon 7 (K205-S206), in 16HBE and GES-1 cell lines.

| Cell line | N | Exon 3 |  | Exon 7 ( KS ) |  | Exon 10 |  |
| --- | --- | --- | --- | --- | --- | --- | --- |
|  |  | n | ratio | n | ratio | n | ratio |
| 16HBE | 29 | 27 | 93.10% | 22 | 75.86% | 19 | 31.03% |
| GES-1 | 27 | 21 | 77.78% | 20 | 74.07% | 12 | 44.44% |

